# ACE2 Pathway Regulates Thermogenesis and Energy Metabolism

**DOI:** 10.1101/2021.08.10.455823

**Authors:** Xi Cao, Tingting Shi, Chuanhai Zhang, Wanzhu Jin, Lini Song, Yichen Zhang, Jingyi Liu, Fangyuan Yang, Charles N Rotimi, Amin Xu, Jinkui Yang

**Author notes:** Address correspondence and reprint requests to Professor JinKui Yang, Department of Endocrinology, Beijing Tongren Hospital, Capital Medical University, Beijing 100730, China. These authors contribute equally to this paper (Xi Cao, Tingting Shi, Chuanhai Zhang).

## Abstract

Identification of key regulators of energy homeostasis holds important therapeutic promise for metabolic disorders, such as obesity and diabetes. ACE2 cleaves angiotensin II (Ang II) to generate Ang-(1-7) which acts mainly through the Mas receptor. Here, we identify ACE2 pathway as a critical regulator in the maintenance of thermogenesis and energy expenditure. We found that ACE2 is highly expressed in brown adipose tissue (BAT) and that cold stimulation increases ACE2 and Ang-(1-7) levels in BAT and serum. *ACE2* knockout mice (*ACE2*^*-/y*^), *Mas* knockout mice (*Mas*^*-/-*^), and the mice transplanted with brown adipose tissue from *Mas*^*-/-*^ mice displayed impaired thermogenesis. In contrast, impaired thermogenesis of db/db obese diabetic mice and high-fat diet-induced obese mice were ameliorated by overexpression of ACE2 or continuous infusion of Ang-(1-7). Activation of ACE2 pathway was associated with improvement of metabolic parameters, including blood glucose, lipids and energy expenditure in multiple animal models. Consistently, ACE2 pathway remarkably enhanced the browning of white adipose tissue. Mechanistically, we showed that ACE2 pathway activated Akt/FoxO1 and PKA pathway, leading to induction of UCP1 and activation of mitochondrial function. Our data propose that adaptive thermogenesis requires regulation of ACE2 pathway and highlight novel therapeutic targets for the treatment of metabolic disorders.

## Introduction

Energy imbalance and the associated metabolic syndromes have become a worldwide public health problem. Thus, identifying factors that can stimulate energy expenditure is instrumental to the development of therapeutics to reduce obesity associated disorders that affect over 10% of the world population (Dong, Lin, Lim, Jin, & Lee, 2017). In the renin-angiotensin system (RAS), angiotensin-converting enzyme 2 (ACE2) cleaves angiotensin II (Ang II) to generate angiotensin-(1-7) (Ang-(1-7)). Ang-(1-7) is a heptapeptide hormone which acts mainly through G-protein-coupled receptor Mas (Santos et al., 2003). ACE2-Ang-(1-7)-Mas pathway works as a negative regulator of ACE-Ang II pathway in multiple disease states (Clarke & Turner, 2012).

In this study we reported the effects of ACE2 pathway on regulating thermogenesis and energy metabolism via modulating mitochondrial function. We found that *ACE2* knockout (*ACE2*^-/y^) and *Mas* knockout (*Mas*^*-/-*^) mice are cold intolerance. We provided compelling genetic, metabolic, physiological, histological, cellular, and molecular evidence to demonstrate that ACE2 pathway is a critical regulator in the maintenance of energy expenditure. This pathway regulates function of brown adipose tissue (BAT) and systemic energy metabolism. Mechanistically, ACE2 pathway activates both Akt/FoxO1 signaling and PKA signaling, leading to induction of uncoupling protein-1 (UCP1) and activation of mitochondrial function. Therefore, ACE2 pathway is a potential treatment target for metabolic disorders including diabetes, obesity, and even cardiovascular diseases.

## Results

### Acute cold exposure increases components of ACE2 pathway

The major tissue of the body where energy is converted into the form of heat to maintain the body temperature is BAT. We found both mRNA level and protein level of ACE2 and Mas in BAT were obviously higher than the ones in subcutaneous white adipose tissue (scWAT) and epididymal white adipose tissue (eWAT) in mice (***Figure 1A and B***). Acute cold exposure caused a significant up-regulation of ACE2 protein expression in BAT (***Figure 1C***). Meanwhile, *ACE2* mRNA levels in BAT, scWAT and eWAT, and *Mas* mRNA levels in BAT and eWAT also increased after exposed to 4°C for 48 hours (***Figure 1D and E***). ACE2 and Ang-(1-7) were also marginally increased in serum upon cold challenge (***Figure 1F and G***). These results demonstrated a selective induction of ACE2 pathway in thermogenic adipose depots (BAT and scWAT) and blood circulation in response to cold environment.

**Figure 1.**
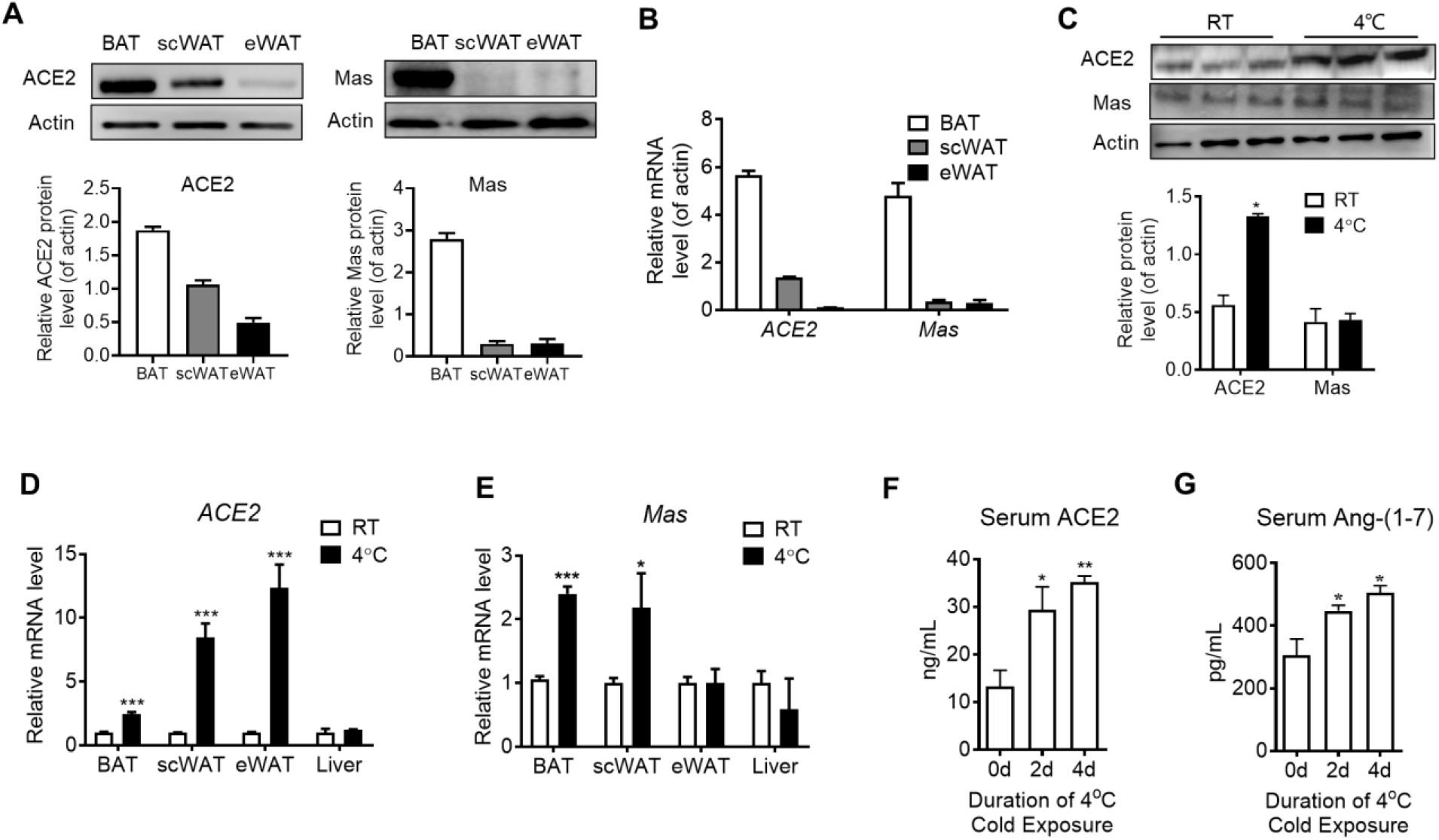
ACE2 pathway is activated by cold exposure. Eight-week-old male C57BL/6J mice were housed at room temperature (RT) for 2 weeks before cold exposure at 4°C for various time periods as indicated. **(A)** Levels of ACE2 and Mas protein from interscapular brown adipose tissue (BAT), subcutaneous and epididymal white adipose tissue (scWAT and eWAT) of C57BL/6 mice at room temperature (RT), as determined by Western blotting. n=3/each group. **(B)** Levels of ACE2 and Mas mRNA from BAT, scWAT and eWAT of C57BL/6 mice at RT, as determined by qPCR. n=5/each group. **(C)** Levels of ACE2 and Mas protein from interscapular BAT of C57BL/6 mice at RT or exposed to 4°C for 6 hours, as determined by Western blotting. n=3/each group. **(D, E)** Levels of ACE2 and Mas mRNA from BAT, scWAT, eWAT and liver of C57BL/6 mice exposed to 4°C for 24 hours, as determined by qPCR. n=5/each group. **(F, G)** Serum levels of ACE2 (**f**) and Ang-(1-7) (**g**), as determined by ELISA. n=6/each group. Data represent mean ± SEM; \**p* < 0.05, \*\**p* < 0.01 and \*\*\**p* < 0.001 *vs* Control group by Student’s *t-*test.

### ACE2 promotes thermogenesis and energy metabolism

To explore the physiological roles of ACE2 in cold-induced adaptive thermogenesis, we used the HFD-induced *ACE2*^-/y^ mice. ACE2 is essential for expression of neutral amino acid transporters in the gut in previous work (Hashimoto et al., 2012). This is consistent with our observation that *ACE2*^-/y^ mice fed an HFD displayed significantly decreased weight compared to wild type (WT) mice (***Figure 2A***). Serum Ang-(1-7) levels were decreased in the *ACE2*^-/y^ mice (***Figure 2B***). Consistent with previous studies (Cao, Yang, Xin, Xie, & Yang, 2014; C. Liu et al., 2012; Niu, Yang, Lin, Ji, & Guo, 2008; Shi et al., 2018; F. Zhang et al., 2016), *ACE2*^-/y^ mice had an impaired glucose tolerance and abnormal lipid profiles (***Figure 2—figure supplement 1A-C***).

**Figure 2.**
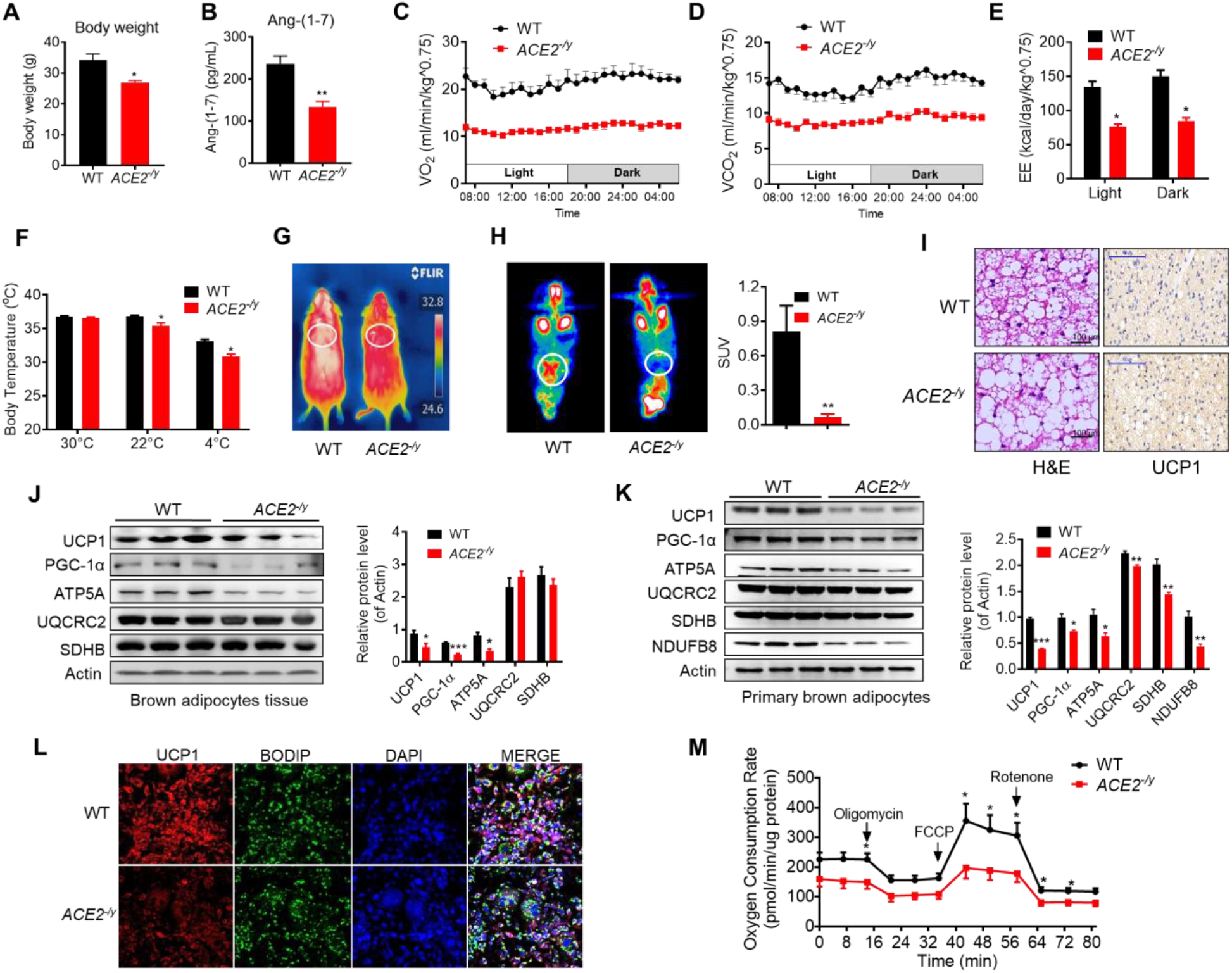
ACE2 deficiency impairs thermogenesis, brown adipose tissue (BAT) activity, and energy metabolism. Eight-week-old male *ACE2*^*-/y*^ mice and their wild type (WT) mice (controls) had a high-fat diet (HFD) for 8 weeks. **(A)** Body weight of *ACE2*^*-/y*^ and WT mice fed a HFD for 8 weeks. **(B)** Serum levels of Ang-(1-7), as determined by ELISA. **(C-E)** Energy expenditure was evaluated by measurement of oxygen consumption (VO_2_) (**C**), carbon dioxide release (VCO_2_) (**D**) and energy expenditure (EE) (**E**) over a 24h period. **(F)** Core body temperature at 30°C, 22°C and 4°C for 8 hours in *ACE2*^*-/y*^ and WT mice. **(G)** Infrared thermal images at 22°C in *ACE2*^*-/y*^ and WT mice. **(H)** Representative tomography–computed tomography (PET-CT) image and standard uptake values (SUVs). **(I)** Representative haematoxylin and eosin (H&E) staining and uncoupling protein-1 (UCP1) immunostaining from BAT sections of *ACE2*^*-/y*^ and WT mice exposure at 4°C. **(J)** Representative western blots showing the changes of key proteins of energy expenditure and thermogenesis in BAT of *ACE2*^*-/y*^ and WT mice exposure at 4°C (n =3/each group). **(K)** Representative western blots showing the key protein changes in primary brown adipocytes from *ACE2*^*-/y*^ and WT mice (n =3/each group). **(L)** Representative immunofluorescent images of *in vitro* differentiated primary brown adipocytes of *ACE2*^*-/y*^ and WT mice, primary brown adipocytes show staining for UCP1 (red), boron-dipyrromethene (BODIPY) (green; neutral lipid dye), and DAPI (blue; nuclei). **(M)** Continuous measurement of oxygen consumption rate (OCR) in primary brown adipocytes from *ACE2*^*-/y*^ mice and WT littermates. Oxygen consumption was performed under basal conditions, following the addition of oligomycin (14μM), the pharmacological uncoupler FCCP (10μM) or the Complex III and I inhibitor antimycin A and rotenone (4μM each). n=5-7/each group unless otherwise stated; Data represent mean ± SEM; \**p* < 0.05, \*\**p* < 0.01 *vs* WT group by Mann-Whitney U test. The online version of this article includes the following figure supplement(s) for figure 2: **Figure supplement 1**. ACE2 deficiency impairs adaptative thermogenesis by cold stimulation.

A key factor for controlling energy homoeostasis is the balance between caloric intake and energy expenditure. Thus, we measured energy expenditure using a comprehensive laboratory animal monitoring system (CLAMS). We observed a decreased oxygen consumption (VO_2_), carbon dioxide release (VCO_2_) and energy expenditure (EE) in *ACE2*^-/y^ mice (***Figure 2C, D and E***), without observable changes in food and/or water intake as well as physical activity, compared to the WT mice (***Figure 2—figure supplement 1D-F***).

To further examine the differences in energy expenditure among these animals, we performed a cold tolerance test in order to gauge adaptive thermogenesis. *ACE2*^-/y^ mice had lower thermogenesis than the WT mice in a cold environment (4°C) (***Figure 2—figure supplement 1G***). To explore the source of thermogenesis, we analyzed the non-shivering thermogenesis (NST) of *ACE2* KO mice in thermoneutral condition (30°C), ambient temperature (22°C) and acute cold (4°C) for 8 hours. *ACE2*^-/y^ mice had lower thermogenesis than the WT mice in either 22 °C or 4°C (***Figure 2F***). This temperature difference was monitored by an infrared camera (***Figure 2G***).

To investigate whether ACE2 induced thermogenesis was related to BAT function, we performed the Positron emission tomography–computed tomography (PET-CT) analysis and the results showed a higher PET-CT signal in BAT of the HFD-induced WT mice than *ACE2*^-/y^ mice (***Figure 2H***). As expected, BAT in *ACE2*^-/y^ mice displayed larger lipid droplets but reduced multilocular structures compared to the WT mice, and had reduced UCP1 expression (***Figure 2I***).

To evaluate the significance of cold-induced ACE2 for thermogenic function of BAT, the expression levels of a network of genes and proteins controlling energy expenditure and thermogenic programming were measured. Protein levels (UCP1, PGC-1α and ATP5A) (***Figure 2J***) and mRNA levels (*UCP1, PRMD16* and PPARγ) (***Figure 2—figure supplement 1H***) in BAT from *ACE2*^-/y^ were obviously decreased.

To validate the above-mentioned change of thermogenesis of BAT was cell autonomous, primary brown adipocytes from *ACE2*^-/y^ mice was fractionated and differentiated *in vitro*. Notably, the protein and mRNA expression of known BAT markers were robustly decreased in *ACE2* deficient primary brown adipocytes (***Figure 2K, Figure 2—figure supplement 1I***). Immunohistochemistry was applied to study the level of UCP1 in primary brown adipocytes differentiated from the BAT of the *ACE2*^-/y^ mice. The result showed the UCP1 expression was reduced in the *ACE2* deficiency primary brown adipocytes (***Figure 2L***). More importantly, the oxygen consumption rate (OCR) was significantly decreased in *ACE2* deficient primary brown adipocytes (***Figure 2M***).

As a complementary approach to the KO mouse models, we carried out gain-of-function studies using ACE2 over expression in obese diabetic db/db mice. One week following adenovirus induced ACE2 over-expression (Ad-ACE2) by tail vein injection in the db/db mice, both ACE2 (***Figure 3—figure supplement 2A***) in BAT and circulating Ang-(1-7) (***Figure 3A***) were increased. Consistent with our previous study, the Ad-ACE2-treated mice exhibited an improved metabolic profile as indicated by the significant alleviation of glucose intolerance (***Figure 3—figure supplement 2B***). Notably, although no observable change on the body weight was observed in the two groups (***Figure 3B***), serum triglyceride levels decreased in the Ad-ACE2-treated mice (***Figure 3—figure supplement 2C***), as well as a minor change in serum cholesterol levels (***Figure 3—figure supplement 2D***).

**Figure 3.**
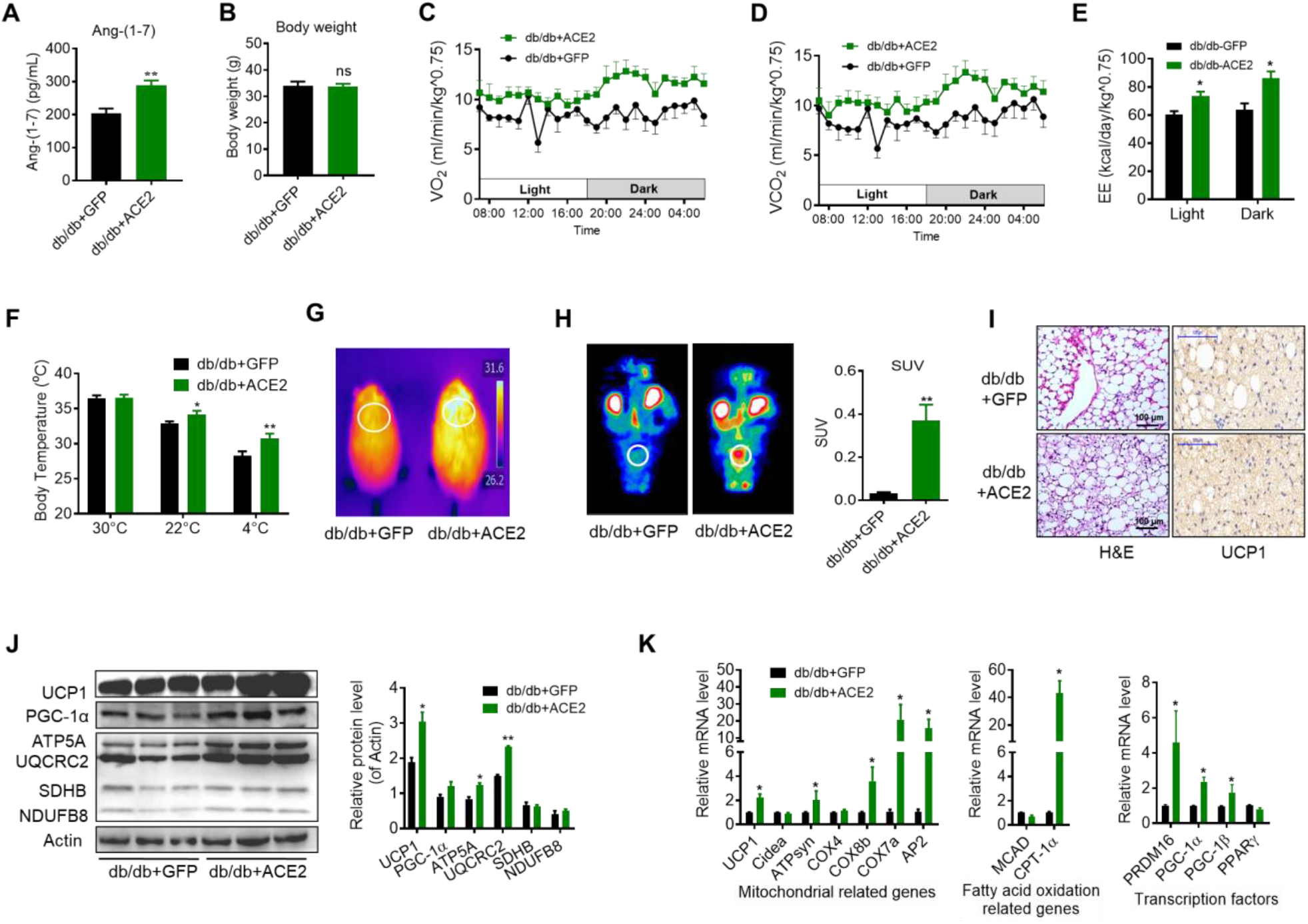
ACE2 enhances thermogenesis, brown adipose tissue (BAT) activity, and energy metabolism in db/db obese mice. ACE2 over-expression adenovirus (Ad-ACE2) and Ad-GFP (control) were introduced into the db/db obese mice by tail vein injection. The ad-ACE2 and Ad-GFP treated db/db mice were used at the 6th day post-virus injection. **(A)** Serum levels of Ang-(1-7), as determined by ELISA **(B)** Body weight of ad-ACE2 and Ad-GFP treated db/db mice at the 6th day post-virus injection. **(C-E)** Energy expenditure was evaluated by measurement of oxygen consumption (VO_2_) (**C**), carbon dioxide release (VCO_2_) (**D**) and energy expenditure (EE) (**E**) over a 24h period. **(F)** Core body temperature at 30°C, 22°C and 4°C for 8 hours. **(G)** Infrared thermal images at 22°C in db/db+ACE2 and db/db+GFP mice. **(H)** Representative tomography–computed tomography (PET-CT) image and standard uptake values (SUVs). **(I)** Representative H&E staining and UCP1 immunostaining from BAT sections of db/db+ACE2 and db/db+GFP mice exposure at 4°C. **(J)** Representative western blots showing the changes of key proteins of energy expenditure and thermogenesis in BAT of db/db+ACE2 and db/db+GFP mice exposure at 4°C. **(K)** Relative mRNA levels of mitochondrial related genes, fatty acid oxidation related genes and transcription factors in BAT of db/db+ACE2 and db/db+GFP mice exposure at 4°C. n=5-7/each group; Data represent mean ± SEM; \**p* < 0.05, \*\**p* < 0.01 *vs* Ad+GFP group by Mann-Whitney U test. The online version of this article includes the following figure supplement(s) for figure 3: **Figure supplement 2**. ACE2 enhance BAT activity and whole-body energy metabolism in db/db mice.

Notably, the Ad-ACE2 treated db/db mice had increased energy expenditure (VO_2_, VCO_2_ and EE) (***Figure 3C-E***). There was no obvious change in food and/or water intake as well as physical activity (***Figure 3—figure supplement 2E-G***).

We measured rectal temperature and infrared thermal imaging in the db/db mice that BAT activity was defective as same as the ones in previous observations (Trayhurn & Wusteman, 1990; Z. Zhang et al., 2014). The results showed that the thermogenesis of the db/db mice was severely impaired (***Figure 3—figure supplement 2H, I***). As expected, the Ad-ACE2 treated db/db mice exhibited better thermogenesis than the control mice in ambient temperature (22°C) and acute cold (4°C) conditions (***Figure 3F, G, Figure 3—figure supplement 2J***). Accordingly, PET-CT result showed that BAT was activated in the Ad-ACE2 treated db/db mice (***Figure 3H***). Moreover, BAT in the Ad-ACE2 treated db/db mice had smaller lipid droplets but increased multi-locular structures, and had increased UCP1 expression compared with the control group (***Figure 3I***).

The protein levels of UCP1, ATP5A and UQCRC2 were significantly increased in the BAT from the Ad-ACE2 treated mice (***Figure 3J***). Consistently, the mRNA levels, including *UCP1, ATPsyn, COX8b, COX7a, AP2, CPT-1α, PRDM16, PGC-1α* and *PGC-1β*, were increased in the BAT from the Ad-ACE2 treated db/db mice (***Figure 3K***). Taken together, these results indicated that ACE2 effectively regulated the mitochondrial biogenesis and respiratory function in brown adipocytes.

### Ang-(1-7) promotes thermogenesis and energy metabolism

To explore the direct physiological roles of Ang-(1-7) in cold-induced adaptive thermogenesis, Ang-(1-7) administration by subcutaneous implantation of micro-osmotic pumps in the db/db and the HFD-induced obese mice were employed. Serum Ang-(1-7) was increased in Ang-(1-7) treated mice (***Figure 4—figure supplement 3A***). There are no significant differences in body weight between the Ang-(1-7) treated db/db mice and the db/db control mice (***Figure 4A***), however, Ang-(1-7) treated db/db mice has an improved glucose tolerance ability (***Figure 4—figure supplement 3B***) and better lipid profiles (***Figure 4—figure supplement 3C, D***). The Ang-(1-7) treated HFD-induced obese mice displayed a lower body weight compared to the control (***Figure 4B***).

**Figure 4.**
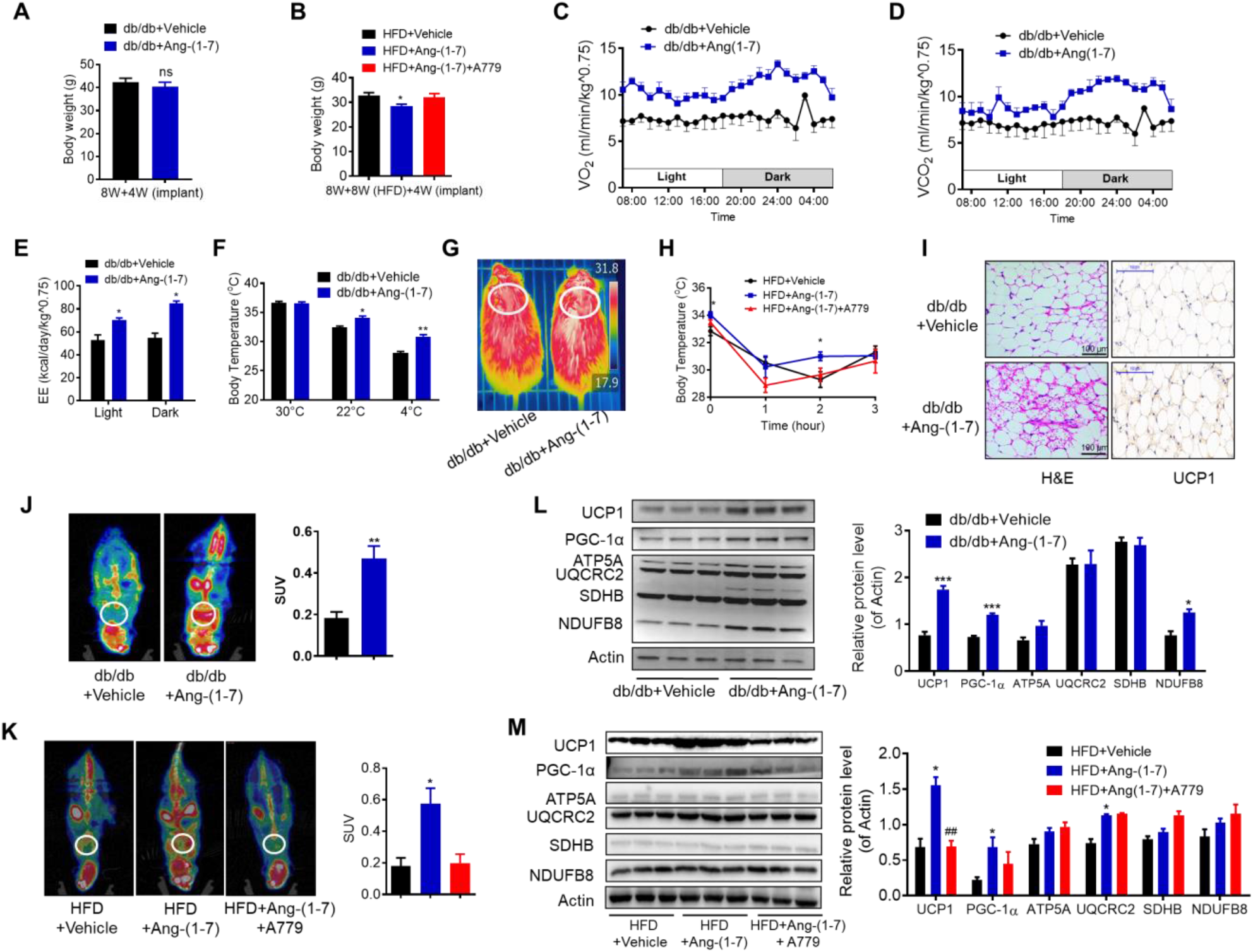
Ang-(1-7) promotes thermogenesis, brown adipose tissue (BAT) activity, and energy metabolism in the db/db and the HFD-induced obese mice. Ang-(1-7) administration by subcutaneous implanted micro-osmotic pumps in the db/db obese mice and the high-fat diet (HFD)-induced obese mice were used. The db/db mice were treated with Ang-(1-7) by subcutaneous infusion of Ang-(1-7) or saline using osmotic mini-pumps for 4 weeks. 6-week-old male C57BL/6J mice were used to develop obesity by HFD diet for 8 weeks, and the mice treated with Ang-(1-7), A779 (an Ang-(1-7) antagonist) or saline by osmotic mini-pumps at the 4th weeks post-HFD. **(A)** Body weight of db/db+Ang-(1-7) and db/db+Vehicle mice at the 4th week post micro-osmotic pumps implantation. **(B)** Body weight of HFD+Ang-(1-7), HFD+A779 and HFD+Vehicle mice at the 4th week post micro-osmotic pumps implantation. **(C-E)** Energy expenditure was evaluated by measurement of oxygen consumption (VO_2_) (**c**), of carbon dioxide release (VCO_2_) (**d**) and of energy expenditure (EE) (**e**) over a 24h period in db/db+Ang-(1-7) and db/db+Vehicle mice. **(F)** Core body temperature at 30°C, 22°C and 4°C for 8 hours in db/db+Ang-(1-7) and db/db-Vehicle mice. **(G)** Infrared thermal images at 22°C in db/db+Ang-(1-7) and db/db+Vehicle mice. **(H)** Core body temperature at 4°C for the indicated lengths of time in HFD+Ang-(1-7), HFD+A779 and HFD+Vehicle mice. **(I)** Representative H&E staining and UCP1 immunostaining from BAT sections of db/db+Ang-(1-7) and db/db+Vehicle mice exposure at 4°C. **(J)** Representative Positron emission tomography–computed tomography (PET-CT) image and SUVs of db/db+Ang-(1-7) and db/db+Vehicle mice. **(K)** Representative PET-CT image and SUVs of HFD+Ang-(1-7), HFD+A779 and HFD+Vehicle mice. **(L)** Representative western blots showing the changes of key proteins of energy expenditure and thermogenesis in BAT of db/db+Ang-(1-7) and db/db+Vehicle mice exposure at 4°C. **(M)** Representative western blots showing the changes of key proteins of energy expenditure and thermogenesis in BAT of HFD+Ang-(1-7), HFD+A779 and HFD+Vehicle mice exposure at 4°C. n=5-7/each group; Data represent mean ± SEM; \**p* < 0.05, \*\**p* < 0.01 *vs* Vehicle group by Student’s *t*-test. The online version of this article includes the following figure supplement(s) for figure 4: **Figure supplement 3**. Ang-(1-7) promoted thermogenesis and energetic metabolism in BAT of db/db mice during cold challenge. **Figure supplement 4**. Ablation of Mas impairs thermogenesis, BAT activity, and energetic metabolism.

Notably, the Ang-(1-7) treated db/db mice had increased energy expenditure (VO_2_, VCO_2_ and EE) (***Figure 4C-E***) without any changes in food and/or water intake as well as physical activity (***Figure 4—figure supplement 3E-G***). Moreover, the Ang-(1-7) treated db/db and the HFD-induced obese mice were better able to defend their body temperature during environmental cold (22°C) and acute cold stress (4°C) compared to the control (***Figure 4F-H, Figure 4—figure supplement 3H***). Meanwhile, the Ang-(1-7) treated db/db mice had increased multi-locular structures but smaller lipid droplets, and increased UCP1 expression comparing to the control group (***Figure 4I***). Accordingly, the Ang-(1-7) treated db/db and the HFD-induced obese mice showed more ^18^F-FDG uptake in the BAT than the control mice recorded by PET-CT (***Figure 4J, K***).

The protein levels of UCP1 and PGC-1α were significantly induced in the BAT from the Ang-(1-7) treated db/db and the HFD-induced obese mice (***Figure 4L, M***). The mRNA levels, including *UCP1, PGC-1α, Cidea, ATPsyn, COX4, COX8b, COX7a, MCAD* and *AP2*, were increased in the BAT from the Ang-(1-7) treated db/db mice (***Figure 4—figure supplement 3I***).

To sum up, our results suggested that the enhanced thermogenesis effect in the Ad-ACE2 and Ang-(1-7) treated mice is caused by the increment of Ang-(1-7) levels, which demonstrates that Ang-(1-7) is crucial to the maintenance of thermogenesis.

### Ablation of Mas impairs thermogenesis in brown adipose tissue

Since the Ang-(1-7), produced by ACE2, realized the function through the Mas receptor, these results above prompted us to hypothesize that the Mas receptor determines the effect of Ang-(1-7) in brown adipose tissue. Firstly, the HFD-induced *Mas*^*-/-*^mice (low Ang-(1-7) action model) were used to assess the therapeutic effects (interventional effects) of Mas on energy metabolism. Although serum Ang-(1-7) levels were increased, the *Mas*^*-/-*^mice had an impaired glucose tolerance, abnormal lipid profiles (***Figure 4—figure supplement 4A-D***), and significantly increased body weight compared to the WT mice (***Figure 4—figure supplement 4E***). Meanwhile, the *Mas*^*-/-*^ mice exhibited decreased oxygen consumption (VO_2_) (***Figure 4—figure supplement 4F***) without any changes in food and/or water intake as well as physical activity (***Figure 4—figure supplement 4G-I***). Moreover, the *Mas*^*-/-*^ mice had lower thermogenesis than the WT mice in either 22 °C or 4 °C (***Figure 4—figure supplement 3J-L***). PET-CT analysis illustrated that the *Mas*^*-/-*^ mice has less ^18^F-FDG uptake in BAT than the WT mice (***Figure 4—figure supplement 4M***). Consistently, the *Mas*^*-/-*^ mice displayed larger lipid droplets and reduced multilocular structures, and had reduced UCP1 expression compared with the WT mice (***Figure 4—figure supplement 4N***). Nevertheless, deletion of *Mas* resulted in a striking repression of BAT thermogenic protein (UCP-1, UQCRC2 and SDHB) (***Figure 4—figure supplement 4O***) and genes (e.g. *UCP1, PRMD16, PGC-1α, PGC-1β, ATPsyn, COX7a* and *CPT-1α*) (***Figure 4—figure supplement 4P***).

To investigate the role of the Mas receptor in BAT, we generated BAT specific *Mas* knockout mice (*Mas*^*-/-*^ BAT transplanted mice). According to the previous studies (X. Liu et al., 2013; Yuan et al., 2016), firstly, BAT of the C57B/L6 recipient mice was removed from the interscapular region. Then, the BAT, which dissected from strain-, sex- and age-matched *Mas*^*-/-*^ donor mice, was subcutaneously transplanted into the dorsal interscapular region of the C57B/L6 recipient mice (WT+*Mas*^*-/-*^-BAT). As controls, C57B/L6 recipient mice transplanted with C57B/L6 BAT (WT+WT-BAT) and C57B/L6 eWAT (WT+WT-eWAT) were used as positive control and negative control, respectively. After the transplantation, the recipient mice were fed by HFD for 10 weeks.

Interestingly, compared with the WT+WT-BAT control mice, the WT+*Mas*^*-/-*^-BAT mice showed greatly impaired HFD-induced insulin resistance. There is no significant difference between the WT+*Mas*^*-/-*^-BAT mice and the WT+WT-eWAT mice in intraperitoneal glucose tolerance test (GTT) and the insulin tolerance test (ITT) (***Figure 4—figure supplement 4Q, R***). Notably, *Mas*^*-/-*^BAT transplantation also strikingly induced HFD-induced weight gain in the WT+*Mas*^*-/-*^-BAT mice compared with the WT+WT-BAT control mice (***Figure 4—figure supplement 4S***).

More importantly, compared to the WT+WT-BAT control mice, the WT+*Mas*^*-/-*^-BAT mice had decreased oxygen consumption (VO_2_), carbon dioxide release (VCO_2_) and energy expenditure (EE) (***Figure 4—figure supplement 4T-V***), along with normal food and/or water intake as well as physical activity (***Figure 4—figure supplement 4W-Y***). Taken together, these results demonstrate that the Mas receptor can directly induce thermogenic program in brown adipose tissues.

### ACE2 pathway induces white fat browning and thermogenesis

Next, we investigated the impact of ACE2/Ang-(1-7) on the process of browning of WAT, a prominent feature in scWAT. Histological examination of scWAT from the Ad-ACE2-treated db/db obese mice showed a profound morphological transformation towards a BAT-like phenotype (smaller adipocytes with multiple lipid droplets) compared with the control (***Figure 5A***). Meanwhile, markers of brown adipocytes, such as UCP1, were significantly increased in scWAT of the Ad-ACE2-treated db/db mice. A much greater induction of transcription factors, including *PRMD16, PGC-1α* and *PGC-1β*, occurred in the scWAT of the Ad-ACE2-treated group (***Figure 5B***). As expected, the Ang-(1-7) treated db/db mice had a similar browning effect as the Ad-ACE2-treated db/db mice in scWAT. Morphological brown-like adipocyte and thermogenic gene expression levels slight increasement were observed in the scWAT after Ang-(1-7) treatment (***Figure 5C, D***).

**Figure 5.**
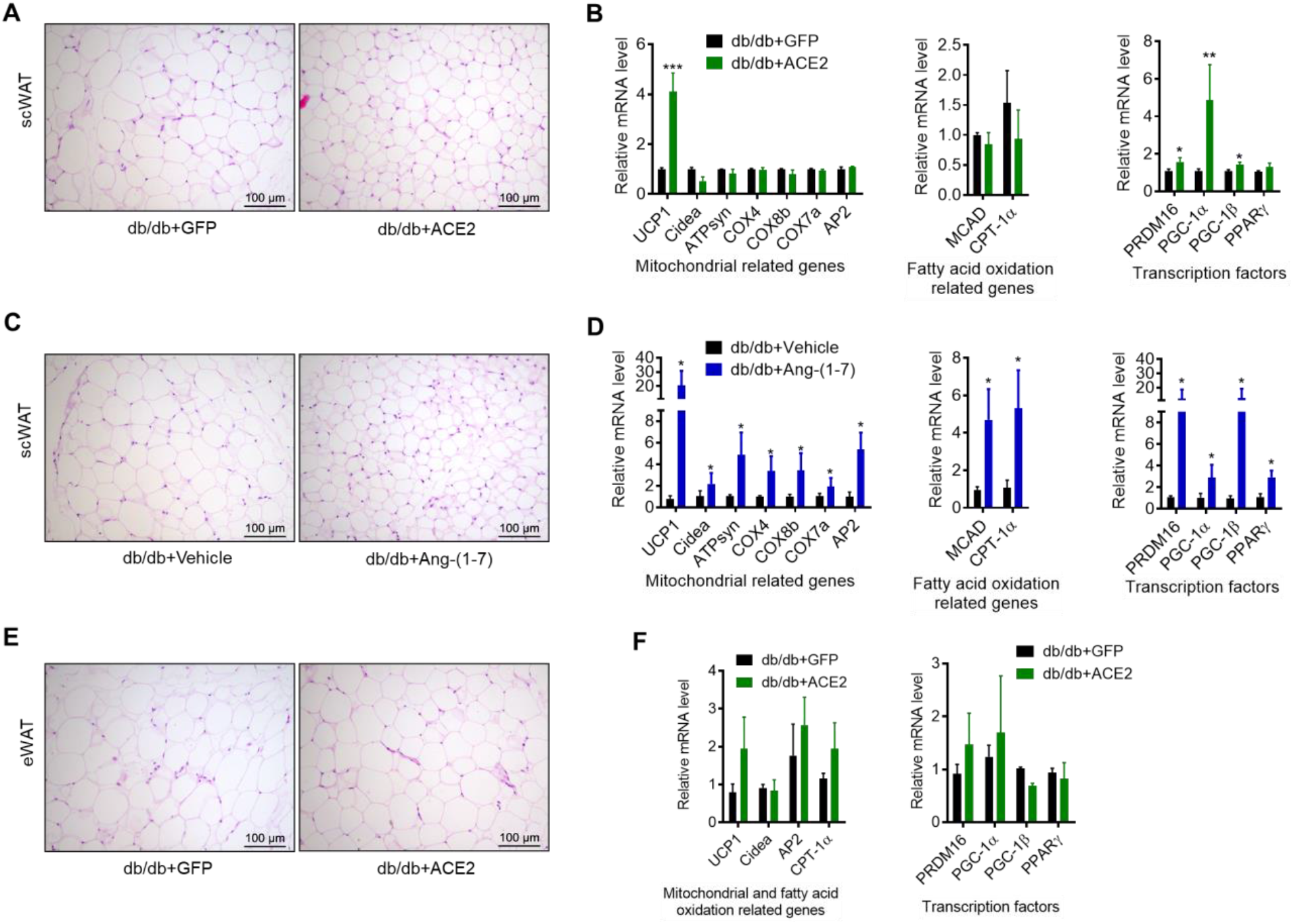
ACE2 pathway induced white adipose tissue browning in the db/db obese mice. **(A)** Representative H&E staining from subcutaneous white adipose tissue (scWAT) sections of db/db+ACE2 and db/db+GFP mice exposure at 4°C. **(B)** Relative mRNA levels of mitochondrial related genes, fatty acid oxidation related genes and transcription factors in scWAT of db/db+ACE2 and db/db+GFP mice exposure at 4°C. **(C)** Representative H&E staining from scWAT sections of db/db+Ang-(1-7) and db/db+Vehicle mice exposure at 4°C. **(D)** Relative mRNA levels of mitochondrial related genes, fatty acid oxidation related genes and transcription factors in scWAT of db/db+Ang-(1-7) and db/db+Vehicle mice exposure at 4°C. **(E)** Representative H&E staining from epididymal white adipose tissue (eWAT) sections of db/db+ACE2 and db/db+GFP mice exposure at 4°C. **(F)** Relative mRNA levels of mitochondrial related genes, fatty acid oxidation related genes and transcription factors in eWAT of db/db+ACE2 and db/db+GFP mice exposure at 4°C. n=5-7/each group; Data represent mean ± SEM; \**p* < 0.05, \*\**p* < 0.01 *vs* GFP/Vehicle group by Student’s *t*-test.

These alterations were restricted to scWAT, but not to eWAT. As shown in Fig. 5e, the morphology and size of eWAT in the Ad-ACE2-treated db/db mice are same compared to the control (***Figure 5E***). Meanwhile, no significant increase in the thermogenic gene expression levels in the eWAT of the Ad-ACE2-treated db/db mice was found (***Figure 5F***).

### ACE2 pathway enhances thermogenesis via Akt and PKA signaling

We further investigated the molecular mechanisms through which ACE2 pathway regulates BAT. Firstly, we performed RNA-seq analysis on BAT isolated from the WT and the *ACE2* KO mice. Notable differences between the two are displayed as 3D-PCA analysis and heat map (***Figure 6—figure supplement 5A, B***). Consistent with the RT-PCR results of BAT in the *ACE2* KO mice, genetic deficiency of *ACE2* significantly altered expression of genes involved in fatty acid biosynthesis, lipid catabolism, lipid biosynthesis, fatty acid beta-oxidation and cholesterol biosynthesis in the BAT of the *ACE2* KO mice (***Figure 6—figure supplement 5C***).

Interestingly, we found that the expression level of Akt associated genes were significantly decreased in the *ACE2* KO mice compared with the WT mice (***Figure 6—figure supplement 5D***). The phosphorylation levels of Akt at residues Thr308 was significantly inhibited in BAT of the *ACE2* KO mice (***Figure 6A***). Furthermore, the phosphorylation levels of Akt were dramatically increased in BAT of the Ad-ACE2 treated mice (***Figure 6B***).

**Figure 6.**
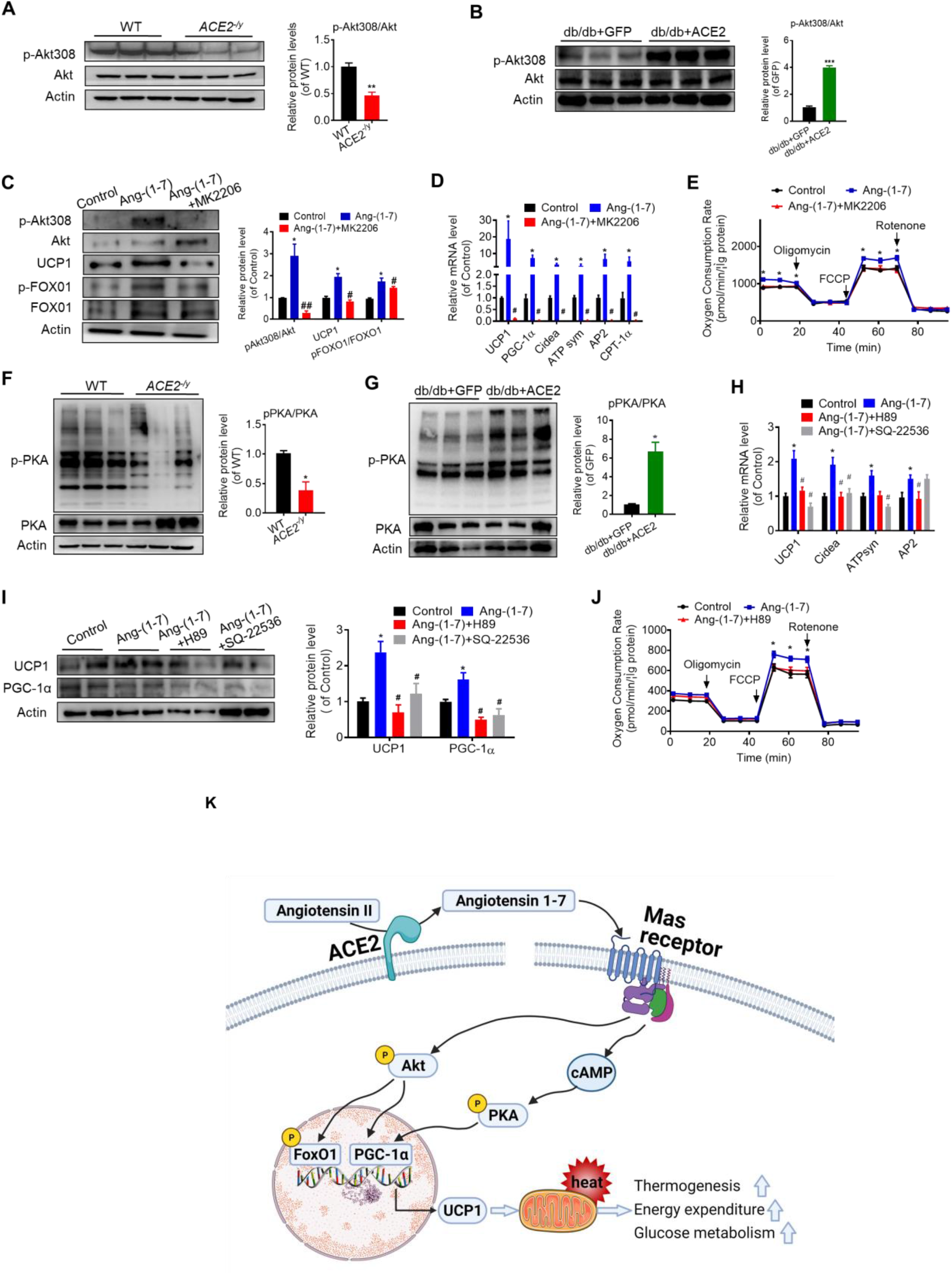
ACE2 pathway induces a thermogenesis program through the Akt signaling and the PKA signaling. Primary brown adipocytes were isolated, cultured, and treated with Ang-(1-7) (10^−6^M) for 24 hours, Akt inhibitor MK2206 (30μM) for 24 hours, PKA inhibitor H89 (30μM) for 2 hours, or adenylylcyclase inhibitor SQ-22536 (10μM) for 24 hours. **(A, B)** Representative western blots showing the changes of p-Akt308 and Akt in BAT of *ACE2*^*-/y*^ (**A**) and db/db+ACE2 mice (**B**) exposure at 4°C. n =3/each group. **(C)** Representative western blots showing the Akt, uncoupling protein-1 (UCP1) and forkhead box protein O1 (FOXO1) changes. n=3/each group. **(D)** Relative mRNA levels of thermogenic and mitochondrial genes. n=4-6/each group. **(E)** Continuous measurement of oxygen consumption rate (OCR). Oxygen consumption was performed under basal conditions, following the addition of oligomycin (14μM), the pharmacological uncoupler FCCP (10μM) or the Complex III and I inhibitor antimycin A and rotenone (4μM each). n =5-7/each group. **(F, G)** Representative western blots showing the p-PKA and PKA changes in BAT of *ACE2*^*-/y*^ (**F**) and db/db+ACE2 mice (**G**) exposure at 4°C. n =3/each group. **(H)** Relative mRNA levels of thermogenic and mitochondrial genes. n=4-6/each group. **(I)** Representative western blots showing the UCP1 and PGC-1α changes. n=4/each group. **(J)** Continuous measurement of OCR. Oxygen consumption was performed under basal conditions, following the addition of oligomycin, the pharmacological uncoupler FCCP (10μM) or the Complex III and I inhibitor antimycin A and rotenone (4μM each). N =5-7/each group. Data represent mean ± SEM; \**p* < 0.05, \*\**p* < 0.01 *vs* control group by Student’s test. # *p* < 0.05, ## *p* < 0.01 *vs* Ang-(1-7) group by Student’s *t*-test. **(K)** Mechanisms involved in ACE2 pathway activation-induced improvement of BAT function. The online version of this article includes the following figure supplement(s) for figure 6: **Figure supplement 5**. RNA-Seq analysis of primary brown adipocytes from *ACE2*^*-/y*^ and WT mice; Ang-(1-7) regulates thermogenesis through Akt and PKA signaling in BAT. **Figure supplement 6**. Ang-(1-7) regulates thermogenesis through Akt and PKA signaling in BAT.

These results prompted us to consider whether ACE2 pathway regulates the function of BAT via Akt signaling. Thus, we treated primary brown adipocytes which isolated from mice with Ang-(1-7). We found phosphorylation of Akt was activated by Ang-(1-7), accompanied by UCP1 up-regulation and forkhead box-containing protein O subfamily-1 (FoxO1) phosphorylation (***Figure 6C***). MK2206, an Akt inhibitor (Chorner & Moorehead, 2018; Matsuzaki, Pouly, & Canis, 2018), suppressed Ang-(1-7) induced UCP1 up-regulation and FoxO1 phosphorylation (***Figure 6C***). Compared to Ang-(1-7) treated primary brown adipocytes cells, MK2206 down-regulated the mRNA levels of *UCP1, PGC-1α, Cidea, ATPsyn, AP2* and *CPT-1α* genes (***Figure 6D***). More importantly, Ang-(1-7) treated primary brown adipocytes exhibited higher OCR, and MK2206 inhibited OCR in the Ang-(1-7) treated primary brown adipocytes (***Figure 6E***). Accordingly, the ACE2-overexpressing primary brown adipocytes cells showed a similar result. MK2206 suppressed ACE2 induced up-regulation of protein (UCP1 and phosphorylated FoxO1) (***Figure 6—figure supplement 6A***) and mRNA *(UCP1* and *CPT-1α*) (***Figure 6—figure supplement 6B***), as well as action on OCR (***Figure 6—figure supplement 6C***). These results suggest that the Akt signaling are required for the thermogenic activity of ACE2 pathway.

After the determination that ACE2 pathway regulates adaptive thermogenesis through the Akt signaling, we paid attention on whether this program could still be provoked by protein kinase A (PKA) signaling, a pathway known to be involved in the canonical thermogenic activation of fat cells. Interestingly, we found that the phosphorylation level of PKA was significantly inhibited in BAT of the *ACE2* KO mice (***Figure 6F***). Similar results appeared in the *Mas* KO mice (***Figure 6—figure supplement 6D***). However, the phosphorylation level of PKA was increased in BAT of the Ad-ACE2-treated db/db mice (***Figure 6G***).

To further elucidate the study on the mechanism of Ang-(1-7)-induced PKA signaling, we treated Ang-(1-7)-treated primary brown adipocytes with PKA and adenylylcyclase inhibitors, simultaneously. Firstly, the Ang-(1-7)-induced PKA signaling was validated in primary brown adipocytes (***Figure 6—figure supplement 6E***). However, H89, a PKA inhibitor, significantly blunted the Ang-(1-7)-induced mRNA levels (*UCP1, Cidea* and *AP2*) (***Figure 6H***) and protein levels (UCP1 and PGC-1α) (***Figure 6I***). Similar effects were observed by using SQ-22536, an adenylylcyclase inhibitor on the formation of intracellular cAMP (***Figure 6H, I***). As expected, the Ang-(1-7) treated primary brown adipocytes exhibited higher OCR, and H89 inhibited OCR in the Ang-(1-7) treated primary brown adipocytes (***Figure 6J***). Our results thus strongly suggest that the PKA signaling is important for the thermogenic activity of ACE2 pathway.

## Discussion

RAS is classically known to regulate blood pressure and maintain water and electrolyte balance. It also plays as a crucial role in metabolic disorders, such as obesity and insulin resistance (Das, 2016). Comprehensive understanding of the complexly biological function of the RAS remains a major biomedical challenge. In the ACE2 pathway, ACE2 is the primary function element, which determines the content of Ang-(1-7), a direct acting peptide, while MAS receptor determines the action form of Ang-(1-7). In order to clarify the role and mechanism of the ACE2 pathway in regulation of thermogenesis, it is necessary to systematically analyze the function of these three elements.

In this study, we used six mice models, *ACE2* KO mice, Ad-ACE2 db/db mice, Ang-(1-7) treated db/db and HFD mice, *Mas* KO mice and BAT-specific *Mas* knockout mice (*Mas*^*-/-*^ BAT transplanted mice), respectively. Based on a series of functional assays in these six mouse models and primary brown adipocytes, we effectively confirmed that the ACE2 pathway regulates glucose and lipid homeostasis. Furthermore, ACE2 pathway maintains thermogenesis and systemic energy metabolism. Molecular analyses, including the use of several inhibitors of Akt and PKA, demonstrate that the effects of ACE2 pathway on brown adipocytes are mediated by both the Akt signaling and the PKA signaling, resulting in the activation of PGC-1α, followed by activation of UCP1 (***Figure 6K***). These findings significantly expand our understanding of the biological function of RAS. Furthermore, our results propose a new concept that the ACE2 pathway can improve obesity and the associated metabolic disorders.

BAT, which utilizes glucose and fatty acids for thermogenesis, contains large number of mitochondria and promotes thermogenesis by mitochondrial respiration through UCP1. BAT-specific UCP1, localized in the inner mitochondrial membrane, plays a fundamental role in thermogenesis. In response to stimulation, activation of PGC-1α up-regulates the expression of BAT-specific UCP1, which dissipates the proton motive force across the inner mitochondrial membranes, and consequentially producing ATP (Sambeat et al., 2016; Schreiber et al., 2017). On the other hand, PGC-1α induces the acquisition of BAT features, including the expression of mitochondria and fatty acid-oxidation and thermogenic genes (Puigserver et al., 1998; Tiraby et al., 2003). We found that the mRNA levels of *UCP1, PGC-1α*, mitochondrial program and fatty acid oxidation related genes (*PGC-1β, ATPsyn, COX7a, COX8b, AP2* and *CPT-1α*) were up-regulated in the *ACE2* overexpression and the Ang-(1-7) treated db/db mice, whereas down-regulated in the *ACE2* KO or the *Mas* KO mice. These results supported that *PGC-1α* and *UCP1* might be critical for the effects of ACE2 pathway on thermogenesis.

We also investigated the underlying mechanisms of ACE2 pathway on the regulation of BAT via the Akt signaling and the PKA signaling (***Figure 6K***).

First, we verified the Akt signaling in the downstream of ACE2 pathway. Akt has a critical function in cell survival and energy balance. Multiple pieces of evidence show that activation of PI3K is followed by the activation of Akt, which in turn triggers a complex cascade of events that include the inhibition of FoxO1 transcription factors and thus the activation of UCP1 and its transcriptional regulator PGC1α (Nakae et al., 2008; Ortega-Molina et al., 2012). In human and 3T3-L1 preadipocytes, Ang-(1-7)-Mas signaling promotes adipogenesis via activation of PI3K-Akt signaling (Than, Leow, & Chen, 2013). AT2R activation induces white adipocyte browning by increasing PPARγ expression, at least in part, via PI3K-Akt signaling pathways (Than et al., 2017). We previously reported that ACE2 and Ang-(1-7) can activate Akt signaling to ameliorate hepatic steatosis (Cao et al., 2016; Cao et al., 2014). In the present study, the *ACE2* KO and the *Mas* KO mice displayed a strong decrease in Akt S308 phosphorylation in BAT. The *ACE2* over-expression or the Ang-(1-7) treatment activated Akt S308 phosphorylation in BAT. Furthermore, the effect of ACE2-Ang-(1-7) on primary brown adipocytes can be attenuated by Akt inhibitor. These results suggest that the Akt signaling might also play a role in ACE2 pathway related regulation of BAT function.

Second, we verified the PKA signaling in the downstream of ACE2 pathway. The Mas receptor was shown to constitutively couple to Gαs, including Gαi, Gαq and Gα12/13proteins (Dias-Peixoto et al., 2008; Gomes, Santos, & Guatimosim, 2012; Tirupula, Desnoyer, Speth, & Karnik, 2014). In the kidney, Ang-(1-7) treatment increased cAMP levels and activated PKA through Gαs activation by the Mas receptor (G. C. Liu, Oudit, Fang, Zhou, & Scholey, 2012; Magaldi, Cesar, de Araujo, Simoes e Silva, & Santos, 2003). Ang-(1-7) regulates insulin secretion through a Mas-dependent cAMP signaling pathway (Sahr et al., 2016). It is well known that norepinephrine released from the sympathetic nerves is a powerful stimulator of BAT. Norepinephrine activates BAT thermogenic program via PKA signaling, followed by the UCP1-mediated proton uncoupling (Su et al., 2017). In this study, the PKA signaling in BAT was changed significantly by ACE2 pathway in mice model. In addition, the effect of ACE2 pathway on primary brown adipocytes can be depressed by cAMP and PKA inhibitor.

Previously, we demonstrate that the ACE2 pathway is involved in the regulation of glucose and lipid homeostasis with limited understanding of the underlying mechanisms (Cao et al., 2014; C. Liu et al., 2012; Niu et al., 2008; Shi et al., 2018; F. Zhang et al., 2016). Here, for the first time, we provide evidence that the alteration in glucose and lipid homeostasis is associated with the change in maintaining brown adipocyte function for the facilitation of energy expenditure. In summary, the ACE2 pathway regulates BAT function and systemic energy metabolisms which is a potential treatment target for metabolic disorders including metabolic syndrome, diabetes, dyslipidemia and fatty liver.

## Materials and methods

### Mice

Obese C57BLKS/J-Leprdb/Leprdb (db/db) male mice, wild-type mice and *Mas* KO mice were purchased from Nanjing Biological Medicine Research Institute, Nanjing University, China. Male C57BL/6J mice were purchased from Vital River Laboratory Animal Technology (Beijing, China). *ACE2* KO mice have been previously described (Niu et al., 2008).

The obese diabetic db/db mice at 7-8 weeks of age were used. Adenovirus (5×10^8^ particle forming units (pfu) in a total volume of 100μL of 0.9% wt/vol saline) was introduced into the db/db mice by tail vein injection. The ad-ACE2 db/db mice were used at the 6th day post-virus injection. The db/db mice were treated with Ang-(1-7) by subcutaneous infusion of Ang-(1-7) (Sigma-Aldrich, St. Louis, MO) (100 ng/kg/min) or saline using osmotic mini-pumps (Alzet-Durect, Cupertino, CA, USA Model #1004) for 4 weeks.

6-week-old male C57BL/6J mice were used to develop obesity by high-fat (HFD) diet (60 kcal% fat) (Research Diets, New Brunswick, NJ, USA) for 8 weeks, and the mice treated with Ang-(1-7) by osmotic mini-pumps at the 5th weeks post-HFD. 8-to 10-weeks old male *ACE2* KO mice and WT controls, *Mas* KO and WT controls were fed HFD diet for 8 weeks before experimental analysis. The mice were housed in a room at controlled temperature (23°C ± 1°C) with a 12-hour light-dark cycle. All animals were handled in accordance with the protocol approved by the Ethics Committee of Animal Research at Beijing Tongren Hospital, Capital Medical University, Beijing, China.

### BAT transplantation

According to the methods described previously (X. Liu et al., 2013; Yuan et al., 2016), BAT was removed from the interscapular region of 8-week old *Mas* KO mice or C57BL/6 mice donor mouse and implanted into the interscapular region of recipient mice. BAT of C57B/L6 recipient mice was removed from the interscapular region. After cervical dislocation of donor mice, the BAT or eWAT (also from the epididymal fat pad of 8-week-old C57BL/6 mice) was removed and peripheral white fat was excluded, and then the remaining BAT (0.2 g) or eWAT (0.2 g) was washed with sterile PBS and transplanted into the interscapular region of recipients as quickly as possible. Recipient mice were anesthetized by ip injection with 400 mg/kg body weight avertin, and then BAT or eWAT was transplanted underneath the skin. The recipient mice were then fed an HFD, which began immediately after the transplantation and continued for 10 weeks.

### Adipocyte oxygen consumption rate (OCR) measurement

Primary brown adipose cells were isolated and cultured for 3 days before plated in XF cell culture microplates (Seahorse Bioscience). Cells (10,000 cells) were seeded in each well and each sample has 8 replicates. After 6 days of differentiation, cultured adipocytes were washed twice and pre-incubated in XF medium for 1-2 h at room temperature. The oxygen consumption rate was measured by the XF extracellular flux analyzer (Seahorse Biosciences). The chemicals (final concentration, 0.2 mM Palmitate: 34 μM BSA, 2 μM Rotenone) were preloaded into cartridges and injected into XF wells in succession. OCR was calculated as a function of time (picomoles per minute).

### Glucose tolerance test (GTT)

Mice were fasted for 16 hours (17:00–9:00) with free access to drinking water. Glucose (1.0 g/kg for the db/db mice and 2.0 g/kg for the HFD mice) was administered intraperitoneally (i.p.), and blood glucose levels were measured immediately 0, 15, 30, 60, and 120 min after glucose injection by using an Accu-Chek glucose monitor (Roche Diagnostics Corp).

### Glucose tolerance test (ITT)

ITT was performed by injecting intraperitoneally 0.75 IU/kg of insulin at mice fasted for 1 hour and measured blood glucose levels at 0, 15, 30, 60, 90, and 120 minutes post injection by using an Accu-Chek glucose monitor (Roche Diagnostics Corp).

### RNA extraction and quantitative Real-time RT-PCR

Total RNA was isolated using TRIzol reagent (Invitrogen, Carlsbad, CA, USA) according to the manufacturer’s instructions. A total of 500ng of RNA was used as the template for the first-strand cDNA synthesis using ReverTra Ace qPCR RT Kit (TOYOBO, Osaka, Japan) in accordance with the manufacturer’s protocol. The transcripts were quantified using Light Cycler 480Real-Time PCR system (Roche, Basel, Switzerland). Primers were designed using Primer Quest (Integrated DNA Technologies, Inc).

### Positron emission tomography–computed tomography (PET-CT)

Siemens Inveon Dedicated PET (dPET) System and Inveon Multimodality (MM) System (CT/SPECT) (Siemens Preclinical Solutions) was used to detect PET-CT imaging at Chinese Academy of Medical Sciences. According to the previously studies (X. Liu et al., 2013; Yuan et al., 2016), mice were allowed to fast overnight and were lightly anesthetized with isoflurane, and then followed by a tail vein injection of 18F-FDG (500 mCi). Sixty mins after the injection of the radiotracer, the mice were subjected to PET/CT analysis. A 10-min CT X-ray for attenuation correction was scanned before PET-CT scan. Static PET-CT scans were acquired for 10 minutes, and images were reconstructed by an OSEM3D algorithm followed by Maximization/Maximum a Posteriori (MAP) or Fast MAP provided by Inveon Acquisition Workplace (IAW) software. The 3D regions of interest (ROIs) were drawn over the guided CT images, and the tracer uptake was measured using Inveon Research Workplace (IRW) (Siemens) software. Individual quantification of the 18F-FDG uptake in each of the ROI was calculated. The data for the accumulation of 18F-FDG on micro PET images were expressed as the standard uptake values (SUVs), which were determined by dividing the relevant ROI concentration by the ratio of the injected activity to the bodyweight. The data are presented as the mean ± SEM.

### Western blot analysis

Tissues were dissolved in RIPA buffer (150mM sodium chloride, 1.0% Triton X-100, 0.5% sodium deoxycholate, 0.1% SDS, 50 mM Tris, protease and phosphatase inhibitor mixture (Roche Diagnostics)). Protein concentrations were determined using a BCA assay kit (Pierce Diagnostics). Protein was separated by 10% (wt/vol) SDS/PAGE, transferred to a PVDF membrane (Millipore), blocked in 5% (wt/vol) skim milk in TBST (0.02 M Trisbase, 0.14 M Vehicle, 0.1% Tween 20, pH 7.4), and incubated with primary antibodies overnight at 4°C and then incubated with secondary antibodies conjugated with HRP. The following primary antibodies were used: anti-UCP1 (ab10983, Abcam), anti-PGC1α (ab54481, Abcam), anti-OXPHOS (ab110413, Abcam), anti-Mas (AAR-013, alomone labs), anti-Akt (#9272, cell signaling technology), anti-p-Akt308 (#13038, cell signaling technology), anti-FOXO1 (#2880, cell signaling technology), anti-p-FOXO1 (#84192, cell signaling technology), anti-PKA (#4782, cell signaling technology), anti-p-PKA (#9621, cell signaling technology), anti-ACE2 (#4335, cell signaling technology), and actin (#4970, Cell Signaling Technology). Signals were detected with Super Signal West Pico Chemiluminescent Substrate (Pierce).

### Histology and immunofluorescence analysis

Tissues fixed in 4% paraformaldehyde were sectioned after being paraffin embedded. Multiple sections were prepared and stained with hematoxylin and eosin for general morphological observations. Cells grown on poly-L-lysine (Sigma)-pretreated coverslips were fixed with 4% paraformaldehyde. Immunofluorescence staining was performed according to the standard protocol using the following antibodies and dilutions: UCP1 (1:100 dilution; Santa Cruz Biotechnologies), MitoTracker Red (1:1000 dilution; Invitrogen). Incubations were performed overnight in a humidified chamber at 4°C. 40, 6-diamidino-2-phenylindole staining was used to mark the cell nuclei. The images were acquired by microscope (DS-RI1; Nikon)

### Metabolic rate and physical activity

Oxygen consumption and physical activity were determined at 12wk of age with a TSE LabMaster, as previously described (Chi & Wang, 2011). Mice were acclimated to the system for 20-24 hours, and then VO_2_ and VCO_2_ were measured during the next 24 hours. Voluntary activity of each mouse was measured with an optical beam technique (Opto-M3, Columbus Instruments, Columbus, OH, USA) over 24 hours and expressed as 24 hours average activity. Heat production and respiratory exchange ratio (RER) were then calculated (X. Liu et al., 2013).

### RNA-Seq analysis

Total RNA was extracted from *ACE2* KO or WT primary brown adipocytes by Trizol reagent (Invitrogen), respectively. Extracted RNA samples were sent to Novel Bioinformatics company (Shanghai, China) for RNA-seq. RNA with RIN>8.0 is right for cDNA library construction. The cDNA libraries were processed for the proton sequencing according to the commercially available protocols. Data were submitted to the GEO archive. Fisher’s exact test was calculated to select the significant pathway, and the threshold of significance was defined by P-value and false discovery rate (FDR) (Dupuy et al., 2007).

### Infrared thermography and core temperature

Mice were exposed to a cold chamber (4°C) with one mouse per cage for up to 6 hours, with free access to food and water. An infrared digital thermographic camera was used to taken images (E60: Compact Infrared Thermal Imaging Camera; FLIR). The images were analyzed by FLIR Quick Report software (FLIR ResearchIR Max 3.4; FLIR). A rectal probe connected to a digital thermometer was used to measure core body temperature (Yellow Spring Instruments).

### Statistical analysis

All of the data are presented as the mean ± SD. The data were analyzed by Student’s t test or one-way ANOVA (with Bonferroni post-hoc tests to compare replicate means) when appropriate. Statistical comparisons were performed using Prism5 (GraphPad Software, San Diego, CA). P values less than 0.05 were considered statistically significant. Representative results from at least three independent experiments are shown unless otherwise stated.

## Acknowledgements

This work was supported by grants from National Key R&D Program of China (2017YFC0909600) and National Natural Science Foundation of China (81561128015, 81471014) to Jinkui Yang, National Natural Science Foundation of China (81670774, 82070850) and Beijing Natural Science Foundation (7162047) to Xi Cao.

## Author contributions

Xi Cao, Tingting Shiand Chuanhai Zhang designed the experiments, performed the experiments and wrote the manuscript. Lini Song, Yichen Zhang, Jingyi Liu and Fangyuan Yang performed the experiments, Wanzhu Jin and Amin Xu designed the experiments. Charles N Rotimi contributed to the interpretation of results and wrote the manuscript. Jinkui Yang conceived the idea for the study, designed the experiments and wrote the manuscript.

## Additional information

### Supplementary materials

Please refer to *Supplementary Figure S1-S6*.

### Data Availability

Sources for materials used in this study are described in Materials and Methods.

## Supplementary Figures

**Figure supplement 1.**
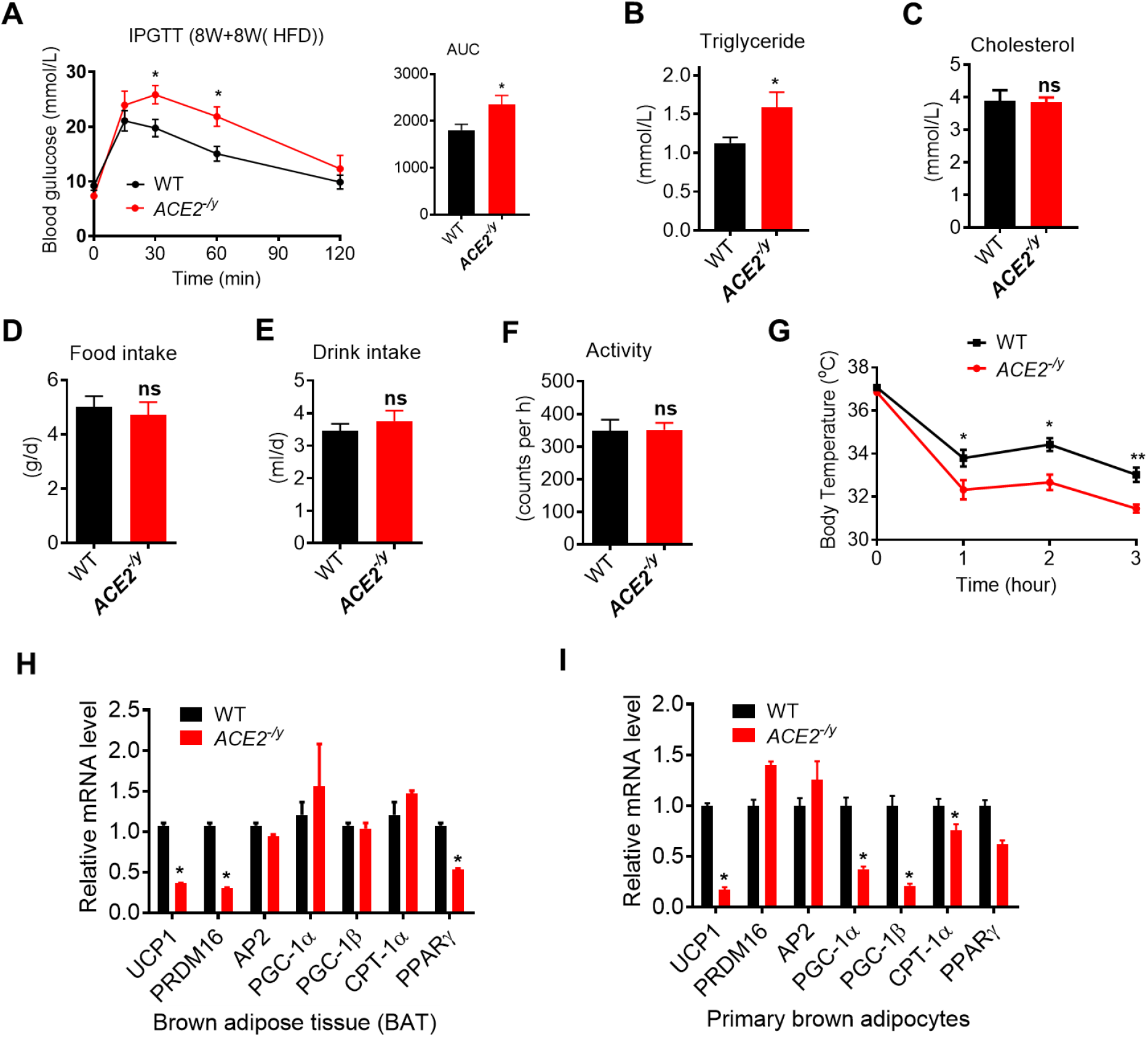
ACE2 deficiency impairs adaptative thermogenesis by cold stimulation. Eight-week-old male *ACE2*^*-/y*^ mice and their WT (control) mice had an HFD for 8 weeks (*ACE2*^*-/y*^ *vs* WT). (**A-C**) Intraperitoneal glucose tolerance test (IPGTT), serum triglyceride and cholesterol levels in *ACE2*^*-/y*^ and WT mice. (**D-F**) 24 h food intake, water intake and physical activity were measured in *ACE2*^*-/y*^ and WT mice. **(G)** Core body temperature at 4°C for the indicated lengths of time in *ACE2*^*-/y*^ and WT mice. **(H)** Relative mRNA levels of mitochondrial related genes, fatty acid oxidation related genes and transcription factors in BAT of in *ACE2*^*-/y*^ and WT mice exposure at 4°C. **(I)** Relative mRNA levels of mitochondrial related genes, fatty acid oxidation related genes and transcription factors in primary brown adipocytes from *ACE2*^*-/y*^ and WT mice. n=5-7/each group; Data represent mean ± SEM; \**p* < 0.05, \*\**p* < 0.01 *vs* WT group by Student’s test.

**Figure supplement 2.**
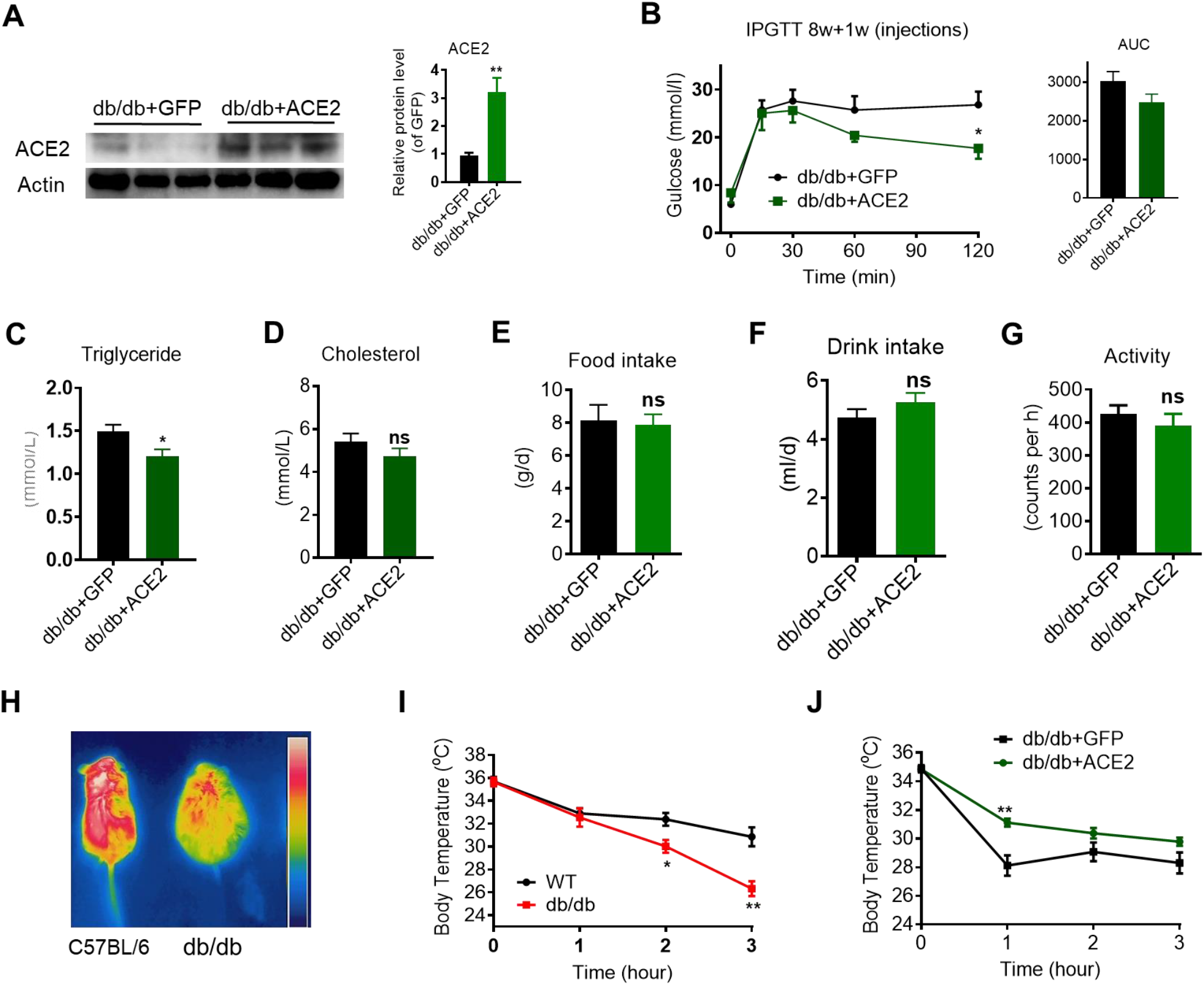
ACE2 enhance BAT activity and whole-body energy metabolism in db/db mice. The ad-ACE2 and Ad-GFP treated db/db mice were used at the 6th day post-virus injection (db/db+ACE2 *vs* db/db+GFP). (**A**) ACE2 overexpression was verified in BAT of Ad-ACE2-treated db/db mice by Western blotting. (**B**-**D**) Intraperitoneal glucose tolerance test (IPGTT), serum triglyceride and cholesterol levels in db/db+ACE2 and db/db+GFP mice. (**E**-**G**) 24 h food intake, water intake and physical activity were measured in db/db+ACE2 and db/db+GFP mice. (**H, I**) Infrared thermal images at 22°C and core body temperature at 4°C for the indicated lengths of time in C57BL/6 and db/db mice. (**J**) Core body temperature at 4°C for the indicated lengths of time in db/db+ACE2 and db/db+GFP mice. n=5-7/each group; Data represent mean ± SEM; \**p* < 0.05, \*\**p* < 0.01 *vs* WT/GFP group by Student’s test.

**Figure supplement 3.**
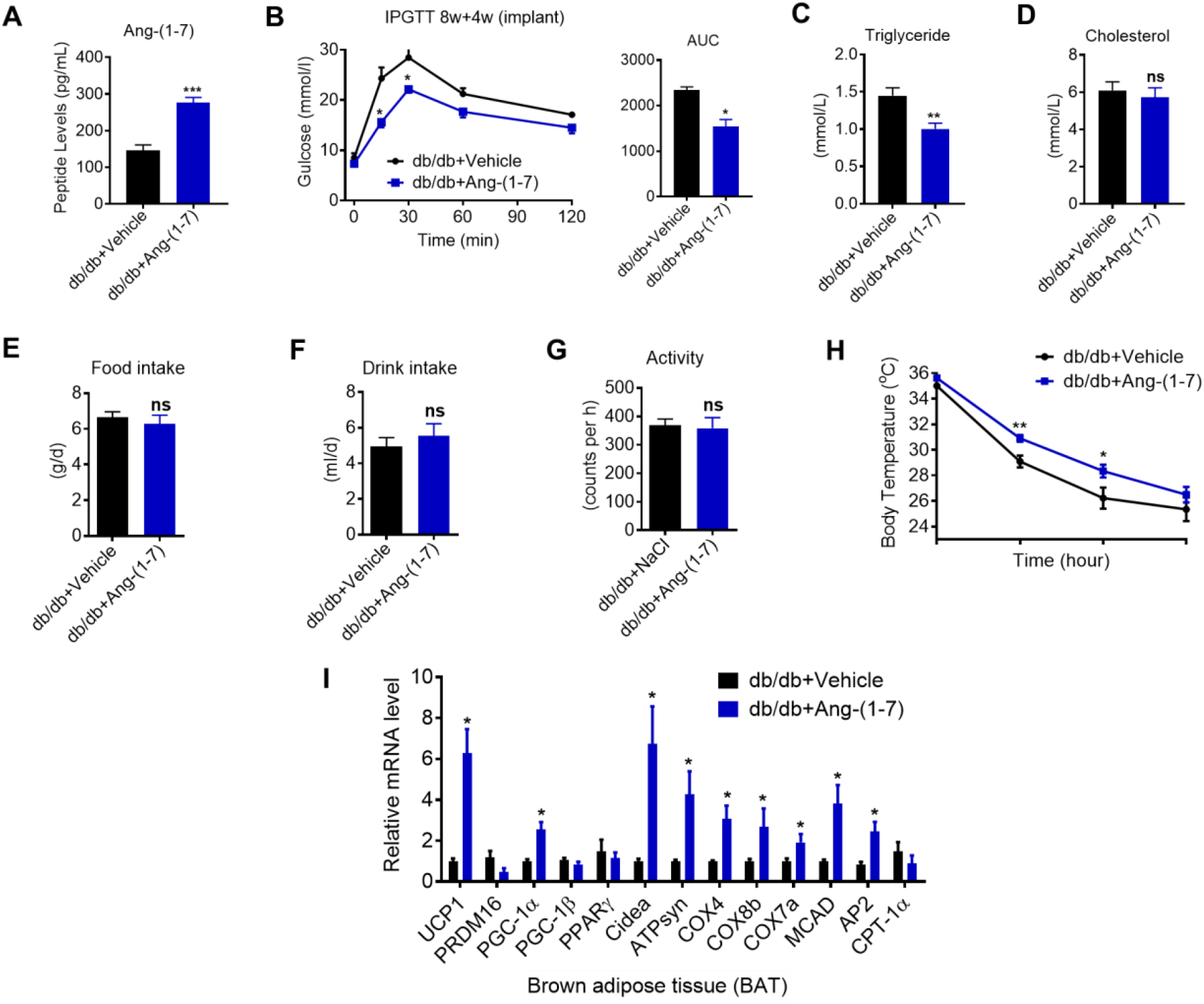
Ang-(1-7) promoted thermogenesis and energetic metabolism in BAT of db/db mice during cold challenge. db/db mice were treated with Ang-(1-7) by subcutaneous infusion of Ang-(1-7) or saline using osmotic mini-pumps for 4 weeks (db/db+Ang-(1-7) *vs* db/db+Vehicle). (**A**) Serum levels of Ang-(1-7) as determined by ELISA in db/db+Ang-(1-7) and db/db+Vehicle mice. (**B**-**D**) Intraperitoneal glucose tolerance test (IPGTT), serum triglyceride and cholesterol levels in db/db+Ang-(1-7) and db/db+Vehicle mice. (**E**-**G**) 24 h food intake, water intake and physical activity were measured in db/db+Ang-(1-7) and db/db+Vehicle mice. **(H)** Core body temperature at 4°C for the indicated lengths of time in db/db+Ang-(1-7) and db/db+Vehicle mice. **(I)** Relative mRNA levels of mitochondrial related genes, fatty acid oxidation related genes and transcription factors in BAT of in db/db+Ang-(1-7) and db/db+Vehicle mice exposure at 4°C. n=5-7/each group; Data represent mean ± SEM; \**p* < 0.05, \*\**p* < 0.01 *vs* Vehicle group by Student’s test.

**Figure supplement 4.**
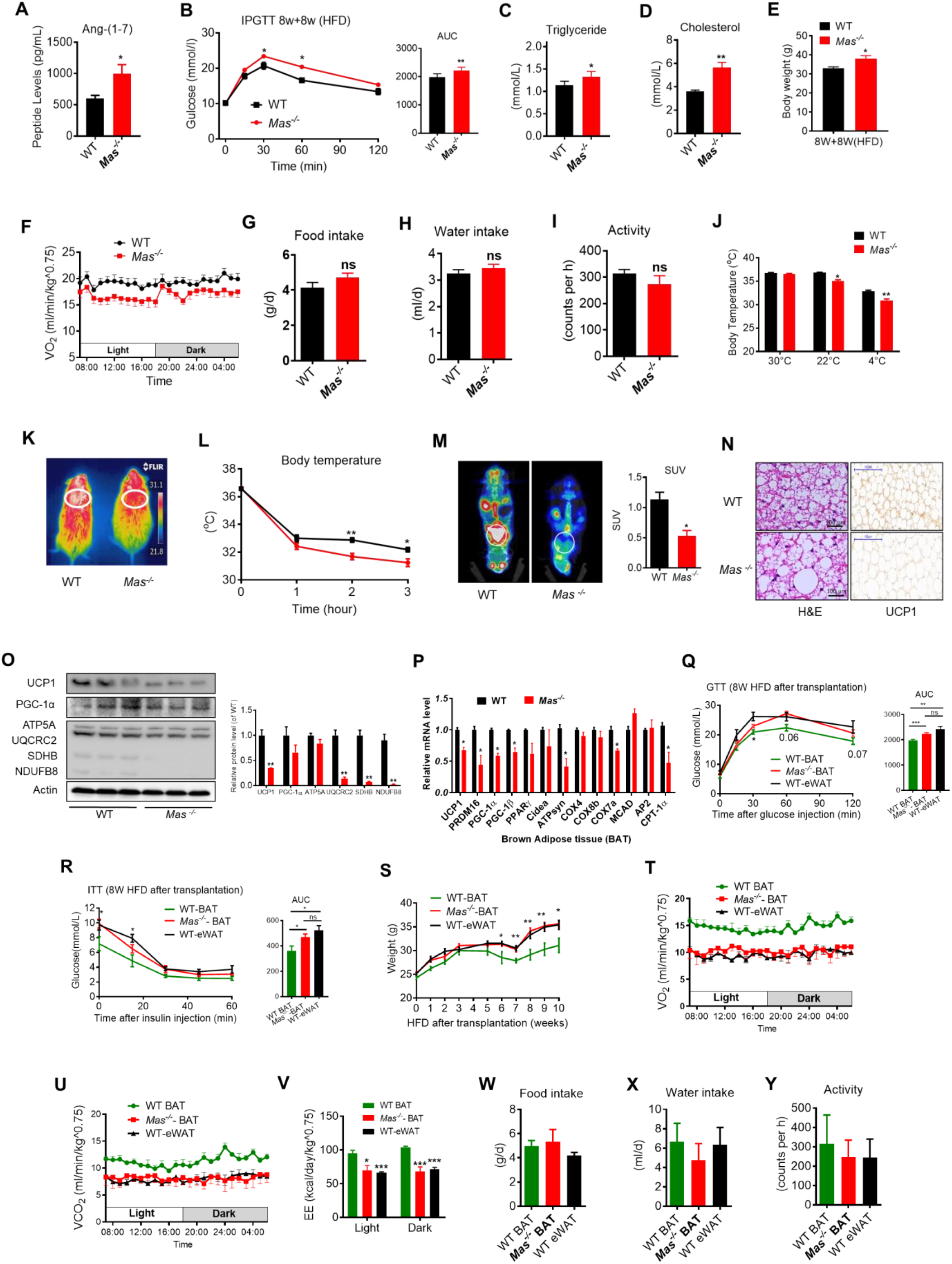
Ablation of Mas impairs thermogenesis, BAT activity, and energetic metabolism. (**A**-**P**) Eight-week-old male *Mas*^*-/-*^ mice and their WT (control) mice had a high-fat diet (HFD) for 8 weeks (*Mas*^*-/-*^ *vs* WT). (**Q**-**Y**) BAT of C57B/L6 recipient mice was removed from the interscapular region. Then, the BAT dissected from *Mas*^*-/-*^ donor mice, was subcutaneously transplanted into the dorsal interscapular region of C57B/L6 recipient mice (WT+*Mas*^*-/-*^-BAT). C57B/L6 recipient mice transplanted with C57B/L6 BAT (WT+WT-BAT) and C57B/L6 epididymal white adipose tissue (eWAT) (WT+WT-eWAT) were used as control. The recipient mice were then fed an HFD immediately after the transplantation and continued for 10 weeks (WT+*Mas*^*-/-*^-BAT *vs* WT+WT-BAT, WT+WT-eWAT). **(A)** Serum levels of Ang-(1-7) as determined by ELISA in *Mas*^*-/-*^ and WT mice. (**B**-**D**) Intraperitoneal glucose tolerance test (IPGTT), serum triglyceride and cholesterol levels in *Mas*^*-/-*^ and WT mice. **(E)** Body weight in *Mas*^*-/-*^ and WT mice fed an HFD for 8 weeks. **(F)** Energy expenditure was evaluated by measurement of oxygen consumption (VO2) over a 24h period in *Mas*^*-/-*^ and WT mice. (**G**-**I**) 24 h food intake, water intake and physical activity were measured in *Mas*^*-/-*^ and WT mice. **(J)** Core body temperature at 30°C, 22°C and 4°C for 8 hours in *Mas*^*-/-*^ and WT mice. **(K)** Infrared thermal images at 22°C in *Mas*^*-/-*^ and WT mice. **(L)** Core body temperature at 4°C for the indicated lengths of time in *Mas*^*-/-*^ and WT mice. **(M)** Representative PET-CT image and SUVs of *Mas*^*-/-*^ and WT mice. **(N)** Representative H&E staining and UCP1 immunostaining from BAT sections of *Mas*^*-/-*^and WT mice exposure at 4°C. **(O)** Representative western blots showing the changes of key proteins in BAT of *Mas*^*-/-*^and WT mice exposure at 4°C. **(P)** Relative mRNA levels of mitochondrial related genes, fatty acid oxidation related genes and transcription factors in BAT of in *Mas*^*-/-*^ and WT mice exposure at 4°C. **(Q)** Intraperitoneal glucose tolerance test (GTT) and the average area under the curve (AUC) in WT+*Mas*^*-/-*^-BAT, WT+WT-BAT and WT+WT-eWAT mice fed an HFD for 8 weeks after transplantation. **(R)** Insulin tolerance test (ITT) and AUC in WT+*Mas*^*-/-*^-BAT, WT+WT-BAT and WT+WT-eWAT mice fed an HFD for 8 weeks after transplantation. **(S)** Body weight time course in WT+*Mas*^*-/-*^-BAT, WT+WT-BAT and WT+WT-eWAT mice fed an HFD over 10 weeks after transplantation. (**T**-**V**) Energy expenditure was evaluated by measurement of oxygen consumption (VO_2_) (**T**), of carbon dioxide release (VCO_2_) (**U**) and of energy expenditure (EE) (**V**) over a 24h period in WT+*Mas*^*-/-*^-BAT, WT+WT-BAT and WT+WT-eWAT mice. (**W**-**Y**) 24 h food intake, water intake and physical activity were measured in WT+*Mas*^*-/-*^-BAT, WT+WT-BAT and WT+WT-eWAT mice fed an HFD for 8 weeks after transplantation. n=5-7/each group; Data represent mean ± SEM; \**p* < 0.05, \*\**p* < 0.01 *vs* WT/ WT+WT-BAT group by Student’s *t*-test.

**Figure supplement 5.**
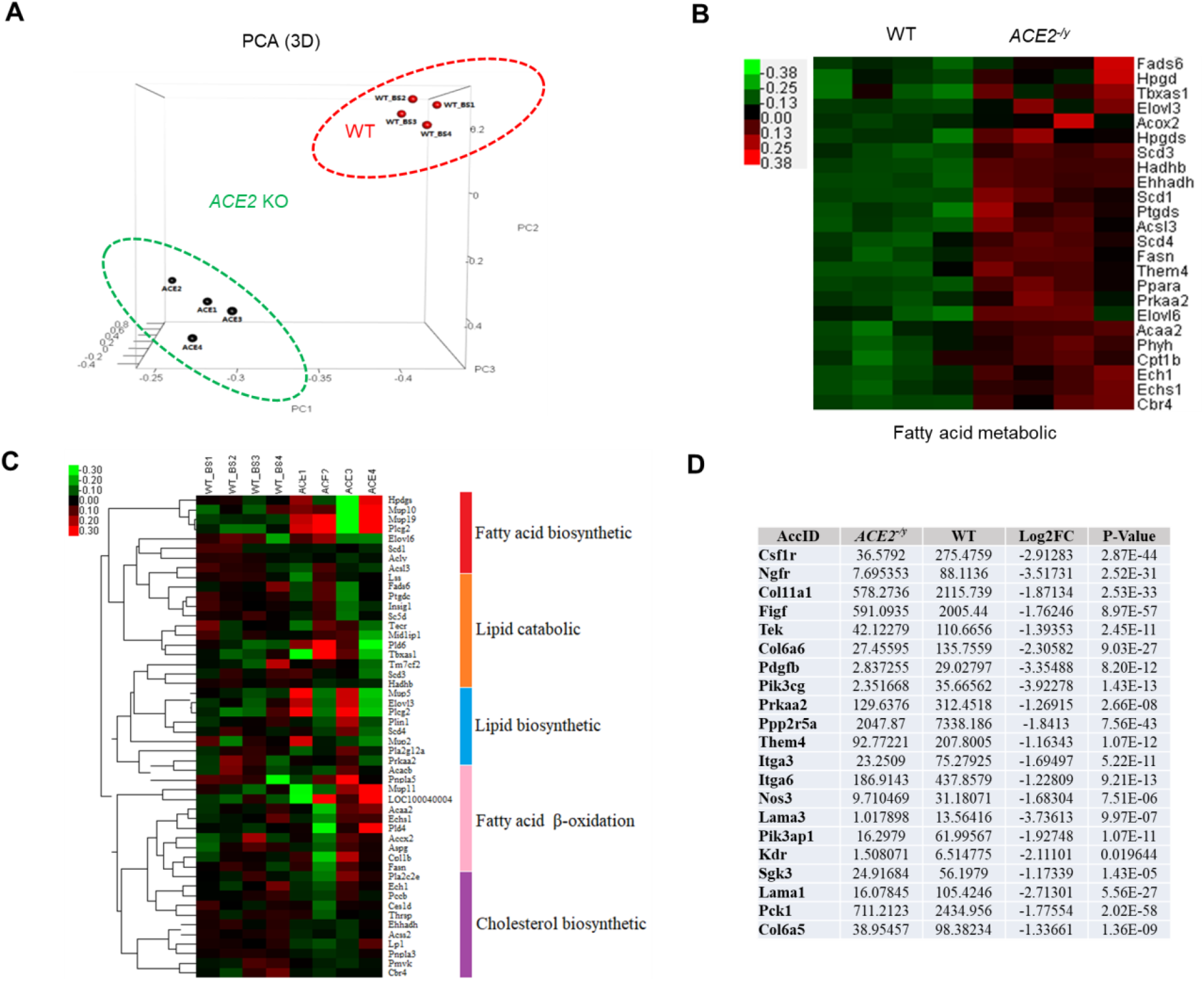
RNA-Seq analysis of primary brown adipocytes from *ACE2*^*-/y*^ and WT mice. **(A)** 3D-PCA analysis represent the deviation of 4 replication within *ACE2*^*-/y*^ (ACE2 1-4) and WT mice (WT_BS 1-4). **(B)** Heat map representation of the differentially expressed genes in *ACE2*^*-/y*^ and WT mice. Gene expression is coded in color scale (−2 to 2). Red or green indicates expression levels above or below the median, respectively. The magnitude of deviation from the median is represented by color saturation. **(C)** Heat map of selected genes associated with fatty acid biosynthetic, lipid catabolic, lipid biosynthetic, fatty acid beta-oxidation and cholesterol biosynthetic, which derived from GO analysis results. **(D)** Table for the expression of Akt signal pathways related genes.

**Figure supplement 6.**
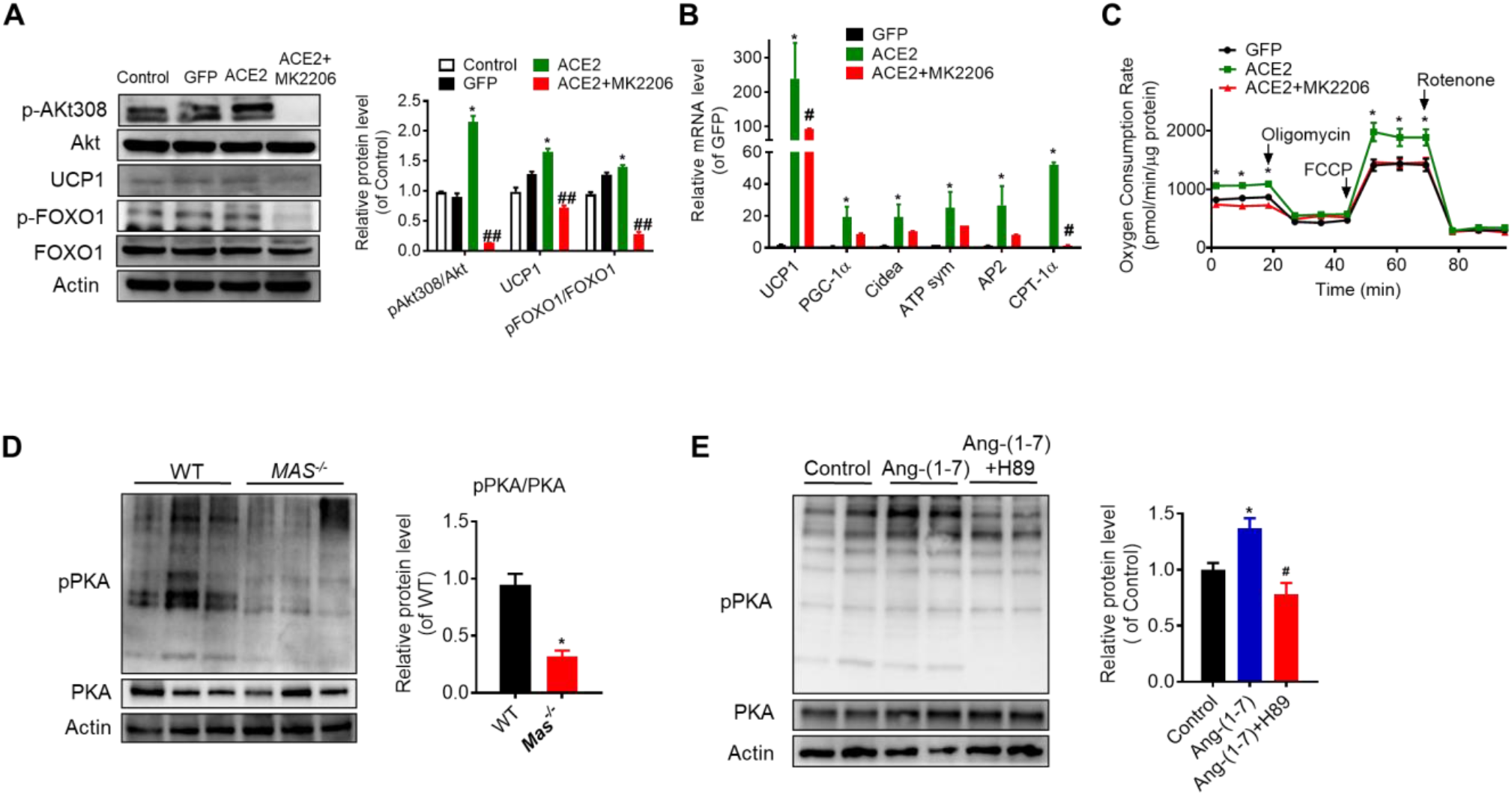
Ang-(1-7) regulates thermogenesis through Akt and PKA signaling in BAT. Primary brown adipocytes were isolated, cultured, infected with ACE2 adenovirus, treated with Ang-(1-7) (10^−6^M) for 24 hours, Akt inhibitor MK2206 (30μM) for 24 hours, or PKA inhibitor H89 (30μM) for 2 hours. **(A)** Representative western blots showing the Akt, UCP1 and FoxO1 changes in primary brown adipocytes overexpressing ACE2 at day 7 and treated with MK2206. n =3/each group. **(B)** Relative mRNA levels of thermogenic and mitochondrial genes. n=4-6/each group. **(C)** Continuous measurement of OCR. Oxygen consumption was performed under basal conditions, following the addition of oligomycin (14μM), the pharmacological uncoupler FCCP (10μM) or the Complex III and I inhibitor antimycin A and rotenone (4μM each). n =4-7/each group. **(D)** Representative western blots showing the p-PKA and PKA changes in BAT of *Mas*^-/-^mice exposure at 4°C. n =3/each group. **(E)** Representative western blots showing the p-PKA and PKA changes in Ang-(1-7) and H89 treated primary brown adipocytes. n =4/each group. Data represent mean ± SEM; \**p* < 0.05, \*\**p* < 0.01 *vs* GFP/WT/control group by Student’s test. # *p* < 0.05, ## *p* < 0.01 *vs* ACE2/Ang-(1-7) group by Student’s *t*-test.

## References

Cao, X., Yang, F., Shi, T., Yuan, M., Xin, Z., Xie, R., Li, S., Li, H., & Yang, J. K. (2016). Angiotensin-converting enzyme 2/angiotensin-(1-7)/Mas axis activates Akt signaling to ameliorate hepatic steatosis. Sci Rep, 6, 21592. doi: 10.1038/srep21592

Cao, X., Yang, F. Y., Xin, Z., Xie, R. R., & Yang, J. K. (2014). The ACE2/Ang-(1-7)/Mas axis can inhibit hepatic insulin resistance. Mol Cell Endocrinol, 393(1-2), 30–38. doi: 10.1016/j.mce.2014.05.024

Chi, Q. S., & Wang, D. H. (2011). Thermal physiology and energetics in male desert hamsters (Phodopus roborovskii) during cold acclimation. J Comp Physiol B, 181(1), 91–103. doi: 10.1007/s00360-010-0506-6

Chorner, P. M., & Moorehead, R. A. (2018). A-674563, a putative AKT1 inhibitor that also suppresses CDK2 activity, inhibits human NSCLC cell growth more effectively than the pan-AKT inhibitor, MK-2206. 13(2), e0193344. doi: 10.1371/journal.pone.0193344

Clarke, N. E., & Turner, A. J. (2012). Angiotensin-converting enzyme 2: the first decade. Int J Hypertens, 2012, 307315. doi: 10.1155/2012/307315

Das, U. N. (2016). Renin-angiotensin-aldosterone system in insulin resistance and metabolic syndrome. J Transl Int Med, 4(2), 66–72. doi: 10.1515/jtim-2016-0022

Dias-Peixoto, M. F., Santos, R. A., Gomes, E. R., Alves, M. N., Almeida, P. W., Greco, L., Rosa, M., Fauler, B., Bader, M., Alenina, N., & Guatimosim, S. (2008). Molecular mechanisms involved in the angiotensin-(1-7)/Mas signaling pathway in cardiomyocytes. Hypertension, 52(3), 542–548. doi: 10.1161/HYPERTENSIONAHA.108.114280

Dong, M., Lin, J., Lim, W., Jin, W., & Lee, H. J. (2017). Role of brown adipose tissue in metabolic syndrome, aging, and cancer cachexia. Front Med. doi: 10.1007/s11684-017-0555-2

Dupuy, D., Bertin, N., Hidalgo, C. A., Venkatesan, K., Tu, D., Lee, D., Rosenberg, J., Svrzikapa, N., Blanc, A., Carnec, A., Carvunis, A. R., Pulak, R., Shingles, J., Reece-Hoyes, J., Hunt-Newbury, R., Viveiros, R., Mohler, W. A., Tasan, M., Roth, F. P., Le Peuch, C., Hope, I. A., Johnsen, R., Moerman, D. G., Barabasi, A. L., Baillie, D., & Vidal, M. (2007). Genome-scale analysis of in vivo spatiotemporal promoter activity in Caenorhabditis elegans. Nat Biotechnol, 25(6), 663–668. doi: 10.1038/nbt1305

Gomes, E. R., Santos, R. A., & Guatimosim, S. (2012). Angiotensin-(1-7)-mediated signaling in cardiomyocytes. Int J Hypertens, 2012, 493129. doi: 10.1155/2012/493129

Hashimoto, T., Perlot, T., Rehman, A., Trichereau, J., Ishiguro, H., Paolino, M., Sigl, V., Hanada, T., Hanada, R., Lipinski, S., Wild, B., Camargo, S. M., Singer, D., Richter, A., Kuba, K., Fukamizu, A., Schreiber, S., Clevers, H., Verrey, F., Rosenstiel, P., & Penninger, J. M. (2012). ACE2 links amino acid malnutrition to microbial ecology and intestinal inflammation. Nature, 487(7408), 477–481. doi: 10.1038/nature11228

Liu, C., Lv, X. H., Li, H. X., Cao, X., Zhang, F., Wang, L., Yu, M., & Yang, J. K. (2012). Angiotensin-(1-7) suppresses oxidative stress and improves glucose uptake via Mas receptor in adipocytes. Acta Diabetol, 49(4), 291–299. doi: 10.1007/s00592-011-0348-z

Liu, G. C., Oudit, G. Y., Fang, F., Zhou, J., & Scholey, J. W. (2012). Angiotensin-(1-7)-induced activation of ERK1/2 is cAMP/protein kinase A-dependent in glomerular mesangial cells. Am J Physiol Renal Physiol, 302(6), F784–790. doi: 10.1152/ajprenal.00455.2011

Liu, X., Zheng, Z., Zhu, X., Meng, M., Li, L., Shen, Y., Chi, Q., Wang, D., Zhang, Z., Li, C., Li, Y., Xue, Y., Speakman, J. R., & Jin, W. (2013). Brown adipose tissue transplantation improves whole-body energy metabolism. Cell Res, 23(6), 851–854. doi: 10.1038/cr.2013.64

Magaldi, A. J., Cesar, K. R., de Araujo, M., Simoese Silva, A.C., & Santos, R. A. (2003). Angiotensin-(1-7) stimulates water transport in rat inner medullary collecting duct: evidence for involvement of vasopressin V2 receptors. Pflugers Arch, 447(2), 223–230. doi: 10.1007/s00424-003-1173-1

Matsuzaki, S., Pouly, J. L., & Canis, M. (2018). In vitro and in vivo effects of MK2206 and chloroquine combination therapy on endometriosis: Autophagy may be required for regrowth of endometriosis. Br J Pharmacol. doi: 10.1111/bph.14170

Nakae, J., Cao, Y., Oki, M., Orba, Y., Sawa, H., Kiyonari, H., Iskandar, K., Suga, K., Lombes, M., & Hayashi, Y. (2008). Forkhead transcription factor FoxO1 in adipose tissue regulates energy storage and expenditure. Diabetes, 57(3), 563–576. doi: 10.2337/db07-0698

Niu, M. J., Yang, J. K., Lin, S. S., Ji, X. J., & Guo, L. M. (2008). Loss of angiotensin-converting enzyme 2 leads to impaired glucose homeostasis in mice. Endocrine, 34(1-3), 56–61. doi: 10.1007/s12020-008-9110-x

Ortega-Molina, A., Efeyan, A., Lopez-Guadamillas, E., Munoz-Martin, M., Gomez-Lopez, G., Canamero, M., Mulero, F., Pastor, J., Martinez, S., Romanos, E., Mar Gonzalez-Barroso, M., Rial, E., Valverde, A. M., Bischoff, J. R., & Serrano, M. (2012). Pten positively regulates brown adipose function, energy expenditure, and longevity. Cell Metab, 15(3), 382–394. doi: 10.1016/j.cmet.2012.02.001

Puigserver, P., Wu, Z., Park, C. W., Graves, R., Wright, M., & Spiegelman, B. M. (1998). A cold-inducible coactivator of nuclear receptors linked to adaptive thermogenesis. Cell, 92(6), 829–839.

Sahr, A., Wolke, C., Maczewsky, J., Krippeit-Drews, P., Tetzner, A., Drews, G., Venz, S., Gurtler, S., van den Brandt, J., Berg, S., Doring, P., Dombrowski, F., Walther, T., & Lendeckel, U. (2016). The Angiotensin-(1-7)/Mas Axis Improves Pancreatic beta-Cell Function in Vitro and in Vivo. Endocrinology, 157(12), 4677–4690. doi: 10.1210/en.2016-1247

Sambeat, A., Gulyaeva, O., Dempersmier, J., Tharp, K. M., Stahl, A., Paul, S. M., & Sul, H. S. (2016). LSD1 Interacts with Zfp516 to Promote UCP1 Transcription and Brown Fat Program. Cell Rep, 15(11), 2536–2549. doi: 10.1016/j.celrep.2016.05.019

Santos, R. A., Simoese Silva, A.C., Maric, C., Silva, D. M., Machado, R. P., de Buhr, I., Heringer-Walther, S., Pinheiro, S. V., Lopes, M. T., Bader, M., Mendes, E. P., Lemos, V. S., Campagnole-Santos, M. J., Schultheiss, H. P., Speth, R., & Walther, T. (2003). Angiotensin-(1-7) is an endogenous ligand for the G protein-coupled receptor Mas. Proc Natl Acad Sci U S A, 100(14), 8258–8263. doi: 10.1073/pnas.1432869100

Schreiber, R., Diwoky, C., Schoiswohl, G., Feiler, U., Wongsiriroj, N., Abdellatif, M., Kolb, D., Hoeks, J., Kershaw, E. E., Sedej, S., Schrauwen, P., Haemmerle, G., & Zechner, R. (2017). Cold-Induced Thermogenesis Depends on ATGL-Mediated Lipolysis in Cardiac Muscle, but Not Brown Adipose Tissue. Cell Metab, 26(5), 753-763.e757. doi: 10.1016/j.cmet.2017.09.004

Shi, T. T., Yang, F. Y., Liu, C., Cao, X., Lu, J., Zhang, X. L., Yuan, M. X., Chen, C., & Yang, J. K. (2018). Angiotensin-converting enzyme 2 regulates mitochondrial function in pancreatic beta-cells. Biochem Biophys Res Commun, 495(1), 860–866. doi: 10.1016/j.bbrc.2017.11.055

Su, J., Wu, W., Huang, S., Xue, R., Wang, Y., Wan, Y., Zhang, L., Qin, L., Zhang, Q., Zhu, X., Zhang, Z., Ye, H., Wu, X., & Li, Y. (2017). PKA-RIIB Deficiency Induces Brown Fatlike Adipocytes in Inguinal WAT and Promotes Energy Expenditure in Male FVB/NJ Mice. Endocrinology, 158(3), 578–591. doi: 10.1210/en.2016-1581

Than, A., Leow, M. K., & Chen, P. (2013). Control of adipogenesis by the autocrine interplays between angiotensin 1-7/Mas receptor and angiotensin II/AT1 receptor signaling pathways. J Biol Chem, 288(22), 15520–15531. doi: 10.1074/jbc.M113.459792

Than, A., Xu, S., Li, R., Leow, M. S., Sun, L., & Chen, P. (2017). Angiotensin type 2 receptor activation promotes browning of white adipose tissue and brown adipogenesis. Signal Transduct Target Ther, 2, 17022. doi: 10.1038/sigtrans.2017.22

Tiraby, C., Tavernier, G., Lefort, C., Larrouy, D., Bouillaud, F., Ricquier, D., & Langin, D. (2003). Acquirement of brown fat cell features by human white adipocytes. J Biol Chem, 278(35), 33370–33376. doi: 10.1074/jbc.M305235200

Tirupula, K. C., Desnoyer, R., Speth, R. C., & Karnik, S. S. (2014). Atypical signaling and functional desensitization response of MAS receptor to peptide ligands. PLoS One, 9(7), e103520. doi: 10.1371/journal.pone.0103520

Trayhurn, P., & Wusteman, M. C. (1990). Lipogenesis in genetically diabetic (db/db) mice: developmental changes in brown adipose tissue, white adipose tissue and the liver. Biochim Biophys Acta, 1047(2), 168–174.

Yuan, X., Hu, T., Zhao, H., Huang, Y., Ye, R., Lin, J., Zhang, C., Zhang, H., Wei, G., Zhou, H., Dong, M., Zhao, J., Wang, H., Liu, Q., Lee, H. J., Jin, W., & Chen, Z. J. (2016). Brown adipose tissue transplantation ameliorates polycystic ovary syndrome. Proc Natl Acad Sci U S A, 113(10), 2708–2713. doi: 10.1073/pnas.1523236113

Zhang, F., Liu, C., Wang, L., Cao, X., Wang, Y. Y., & Yang, J. K. (2016). Antioxidant effect of angiotensin (17) in the protection of pancreatic beta cell function. Mol Med Rep, 14(3), 1963–1969. doi: 10.3892/mmr.2016.5514

Zhang, Z., Zhang, H., Li, B., Meng, X., Wang, J., Zhang, Y., Yao, S., Ma, Q., Jin, L., Yang, J., Wang, W., & Ning, G. (2014). Berberine activates thermogenesis in white and brown adipose tissue. Nat Commun, 5, 5493. doi: 10.1038/ncomms6493

